# Behavioural and neural mechanisms for stochastic choices in mixed-strategy games

**DOI:** 10.64898/2026.07.29.741515

**Authors:** Joanna Aloor, Timothy P.H. Sit, Oliver M. Gauld, Joseph Warren, Matthew Mower, Daeyeol Lee, Chunyu A. Duan

**Author notes:** Authors contributed equally to the work (Joanna Aloor, Timothy P.H. Sit).

## Abstract

Adaptive behaviour usually requires exploiting regularities in the environment, but in competitive settings the opposite can be true: predictable choice patterns can be exploited by others, making unpredictability itself advantageous. How neural circuits generate such strategic variability remains poorly understood. Here, we trained mice to play a zero-sum game against an opponent that exploited statistical regularities in their choices and rewards, and tracked their behaviour and dorsal cortical dynamics across learning. Using a hidden Markov model, we found that mice transitioned from structured, predictable strategies towards a near-optimal stochastic strategy as they learned. Applying the same framework to monkeys playing the same game identified a shared stochastic strategy across species, despite differences in how it was deployed. Cortex-wide imaging revealed that stochastic choices were associated with reduced representation of reward history, while immediate reward signals remained robust. Critically, while the strength of cortical reward signals predicted subsequent choice during reward-guided behaviour, this relationship was abolished during stochastic behaviour. Thus, adaptive stochasticity does not simply arise from a loss of reward information, but from selectively decoupling reward from future choice. These results reveal a neural mechanism through which animals suppress otherwise useful reward-guided structure to generate adaptive unpredictability in competitive environments.

## Introduction

In nature, unpredictable and stochastic behaviour is crucial for survival, particularly in situations where individuals compete with adaptive opponents to maximise reward (Giraldeau and Caraco, 2000; Humphries and Driver, 1970; M. Flaxman, 2000). Predator-prey interactions, territorial disputes, and competition for food or mates all require animals to continuously balance exploration and exploitation whilst avoiding patterns that can be exploited by others. Similarly, humans rely on strategic variability when navigating uncertain social and economic environments, including negotiations, competitive markets, and interactions in which the actions of others influence future outcomes. Understanding how the brain generates flexible and unpredictable choices in multi-agent contexts is therefore central to understanding adaptive decision-making.

Competitive games provide a powerful framework for studying this problem (Camerer, 2003). Game-theoretic analysis predicts that optimal behaviour in such settings corresponds to a mixed-strategy Nash equilibrium, in which actions are selected probabilistically such that no player can improve their expected outcome by unilaterally changing their strategy (Nash, 1950; von Neumann and Morgenstern, 1944). Previous studies have shown that humans, monkeys, and rodents learn to approach a mixed-strategy when playing competitive games (Barraclough, Conroy, and Lee, 2004; Dorris and Glimcher, 2004; Philippe et al., 2024; Tervo et al., 2014; Vickery, Chun, and Lee, 2011; Wang and Kwan, 2023). Different hypotheses have been proposed to describe how animals learn this behaviour. One explanation is that choice stochasticity is driven by model-free reinforcement learning where choice values converge to be equal (Lee et al., 2004). An alternative hypothesis is that animals adopt a counter-predictive strategy, where choices are guided by the opponent’s history of past choices (Barraclough, Conroy, and Lee, 2004; Tervo et al., 2014). However, these models assume animals use a single, stationary decision-making process, whereas recent studies have demonstrated that rodents and other invertebrates often switch between distinct behavioural modes across both short and long timescales (Ashwood et al., 2022; Calhoun, Pillow, and Murthy, 2019). Whether state-switching approaches can explain game-theoretic strategies, and the extent to which identified states generalise across species, remain an open question. State-based behavioural models provide a framework to identify transitions between latent strategies within individual sessions, overcoming limitations of session-averaged measures that obscure dynamic changes in behaviour. This enables more direct comparisons of behavioural and neural representations across distinct behavioural states, providing the means to identify the underlying mechanisms driving stochastic choices.

Competitive decision-making tasks where subjects are required to be unpredictable have revealed that strategic behaviour is supported by distributed frontal cortical representations of reward and choice history, action value, and hypothetical outcomes (Abe and Lee, 2011; Barraclough, Conroy, and Lee, 2004; Lee and Seo, 2007; Seo and Lee, 2007; Seo et al., 2014; Tervo et al., 2014). Complementary mesoscale imaging approaches have further shown that cortical networks encode additional decision variables, including abstract choices and motor actions, across widespread regions (Gauld et al., 2026; Musall et al., 2019; Orsolic et al., 2021; Pinto et al., 2019; Steinmetz et al., 2019). This highlights that cortical circuits maintain rich representations of information relevant for guiding decisions, including signals related to previous outcomes and actions. Yet, when competing against adaptive opponents, reliance on recent reward history undermines performance by introducing predictable biases in behaviour. It remains unclear how neural circuits selectively modulate the influence of outcome information to support stochastic choice while maintaining rich representations of decision-relevant variables. Understanding how cortical circuits balance these competing demands is critical for explaining how animals generate adaptive strategies in uncertain and competitive environments.

Here, we probed how head-fixed mice used stochastic strategies in matching pennies, a two-choice competitive game played against a computer opponent. We used an unsupervised hidden Markov model to characterise how mice switched between stochastic and non-stochastic behavioural states across learning, and demonstrated that this model reproduced findings from a previous monkey dataset. We found that both mice and monkeys flexibly adapted their strategies against different computer opponents. In addition, both species shared a reward-maximising, stochastic behavioural state when needed, while their engagement with stochastic and non-stochastic strategies within sessions differed. To investigate the underlying neural mechanisms, we used widefield calcium imaging and tracked dorsal cortical activity in mice throughout learning. We found that reward history representation was reduced during stochastic choices, whilst immediate reward information remained strongly represented but dissociated from influencing choice switching. These findings support a putative cortical mechanism for stochastic choices through the decoupling of immediate rewards from upcoming choice.

## Results

### Mice learned to make unpredictable choices against an exploitative computer opponent

We trained head-fixed mice to play a matching pennies game against an exploitative computer opponent, where optimal behaviour was to choose randomly (Figure 1a). Following an auditory cue indicating the start of the trial, two lick ports extended within reach of the animal, prompting the beginning of the choice window (Figure 1b). Mice licked left or right to indicate their choice and were rewarded only if their choice differed from the computer’s prediction. The computer opponent used 1-4 trial patterns in the animal’s choice and reward history to predict their upcoming choice, similar to previous opponent designs (Algorithm 2 in Barraclough, Conroy, and Lee, 2004, Competitor 1 in Tervo et al., 2014, and Wang et al., 2022, described in Methods). On each trial, the computer performed the binomial tests for the conditional probabilities for a left choice following the previous 1-4 trial-back patterns; the most significant binomial test determined the computer’s prediction. Therefore, the animal was only rewarded if their choice remained unpredictable to the computer opponent, for which the optimal strategy achieves a 50% reward rate.

**Figure 1.**
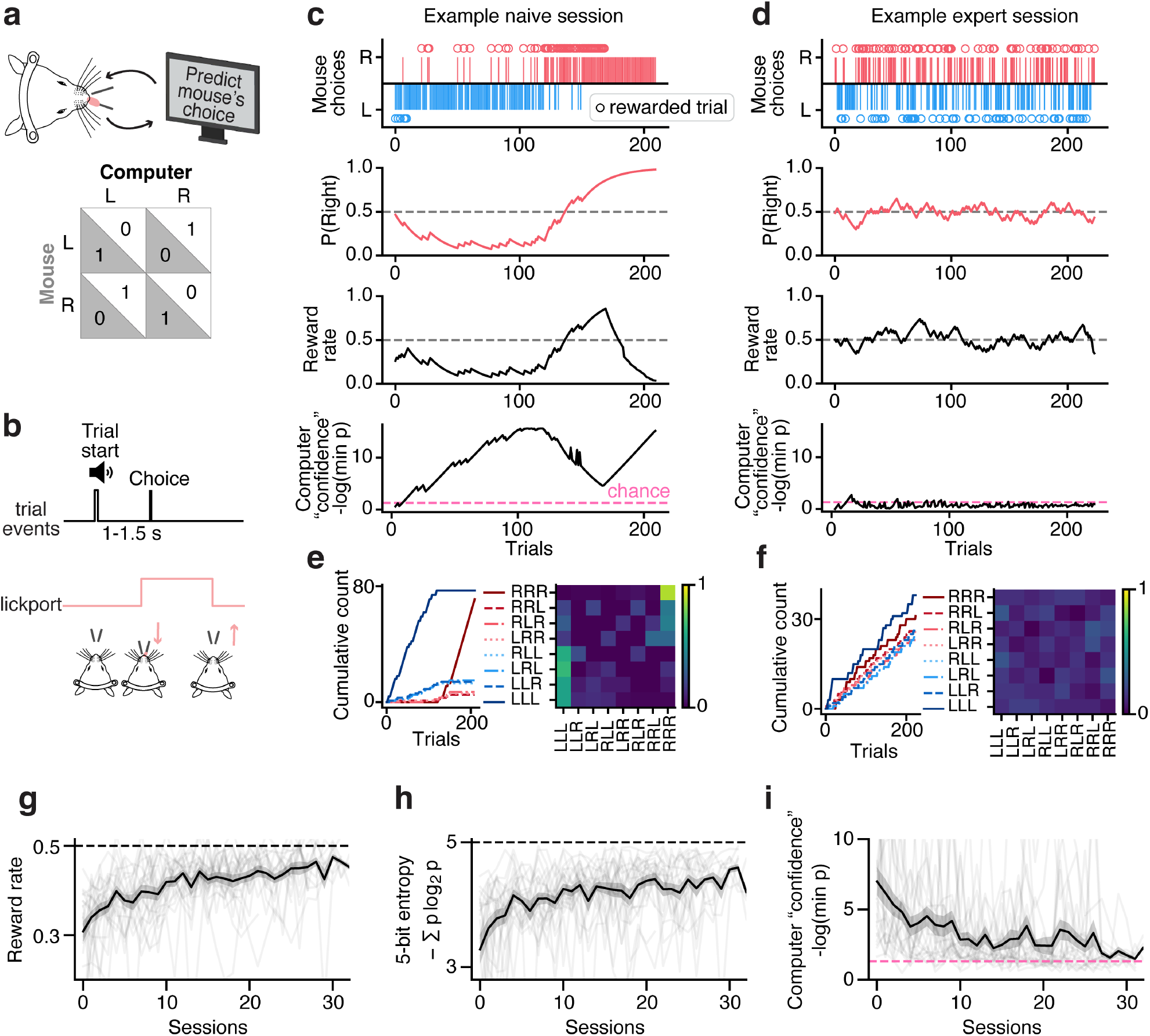
Head-fixed mice learn to make unpredictable decisions against an exploitative computer opponent. **a**, Matching pennies game set-up. On each trial, a head-fixed mouse makes a left or right choice and is rewarded if computer opponent does not correctly predict choice. **b**, Example trial structure. **c**, Example session of a naive mouse playing matching pennies. Top row, right (red) and left (blue) mouse choices, circles indicate rewarded trials. Second row, smoothed probability of right choice. Third row, smoothed reward rate. Bottom row, negative log of the minimum p-value associated with the binomial tests of the occurrences of 1-4 length patterns in the mouse’s full choice and reward history. This can be used as a proxy for computer “confidence” in its predictions. Dashed pink line indicates where p = 0.05, the threshold distinguishing random prediction (p > 0.05) from predictions based on the animal’s recent four-choice-and-reward history (p < 0.05). **d**, Same as (c) for an example expert session from the same mouse. **e**, Left, cumulative count of all 2^3^ possible three-choice patterns for the same example naive session as (c). Right, transition matrices showing the frequency of transitions between all 3-choice patterns, calculated using three-trial sequences while shifting the window by one trial. **f**, Same as (e) for the example expert session in (d). **g-i**, Reward rate (**g**), three-choice entropy (**h**), and median computer “confidence” (i) across learning (n = 22 mice). Dashed pink line indicates where p = 0.05.

Early in training, mice used highly structured choice patterns, often with strong side biases for extended runs of trials, resulting in suboptimal reward rates falling below 0.5 (Figure 1c). Over a few weeks of gameplay, mice learned to generate more stochastic choices that achieved a consistent reward rate close to the optimum (Figure 1d). To better evaluate the animals’ randomness in choices, we utilised the computer opponent’s prediction system that intrinsically tested the animals’ trial-by-trial predictability against the computer’s algorithm. We calculated the negative log of the most significant p-value associated with the computer’s statistical tests to approximate the computer’s “confidence” in predicting the animal’s upcoming choice: larger values indicated more predictable animal choices. In the example early session where the animal made more structured choices, the computer “confidence” remained high throughout the session (Figure 1c bottom). Whereas in the expert session, the computer “confidence” decreased, often falling below the significance threshold (p=0.05), at which point the opponent’s predictions were selected at random (Figure 1d bottom). Consistent with this, animals had highly skewed choice patterns and transition structure during early sessions; whereas in expert sessions, all choice patterns were represented and transitions between patterns were more equally distributed (Figure 1e-f). We found that as the animals’ performance increased, their behavioural entropy significantly increased (5-bit entropy=3.73 ± 0.48 to 4.38 ± 0.31; paired t-test; T = -6.89, p = 8.3 × 10^−7^; n = 22 mice), and the median computer “confidence” decreased (5.42 ± 2.58 to 2.38 ± 1.89; T = 5.44, p = 2.1 × 10^−5^; n = 22 mice; 29 ± 1.1 sessions; Figure 1g-i). Overall, the expert session more closely matched the behaviour of a true random agent choosing from a Bernoulli(0.5) distribution (Extended Figure 1). These results indicate that mice learned to make increasingly stochastic choices against an exploitative computer opponent (Wang et al., 2022).

Although animals increased their general choice stochasticity across learning, sessions often included a mixture of both predictable and unpredictable blocks of trials, suggesting that session-level metrics alone cannot fully distinguish between changes in strategies that animals may be using. This motivated a more detailed characterisation of trial-by-trial behavioural strategy, with the prediction that a reward-maximising, stochastic strategy would emerge as a dominant state in expert animals.

### GLM-HMM reveals transitions from biased to stochastic strategies across learning

To identify discrete latent states underlying trial-by-trial fluctuations in choice behaviour, we fit a hidden Markov model in which each latent state was defined by a generalised linear model (GLM-HMM) mapping trial history covariates to choice probability (Figure 2a; Ashwood et al., 2022). The GLM-HMM has primarily been applied to perceptual tasks to separate engaged from disengaged states or to recover differences in perceptual sensitivity, but its structure is well suited for uncovering latent strategies in competitive games. To predict left and right choice probabilities, we used a GLM with 3 input regressors: general side bias, repeating the previous rewarded choice (repeat-win; 1 for right choice, -1 for left choice), and repeating the previous unrewarded choice (repeat-loss). We validated our model using synthetic simulations of four different strategies: side bias, perseverance by repeating the previous choice, win-stay/lose-switch (WSLS), and random choices. We fit the simulated choices using the GLM-HMM and verified that our approach could recover the ground-truth parameters from a multi-strategy agent that switched between the four predefined strategies according to a known transition matrix (Figure 2b, Extended Figure 2).

**Figure 2.**
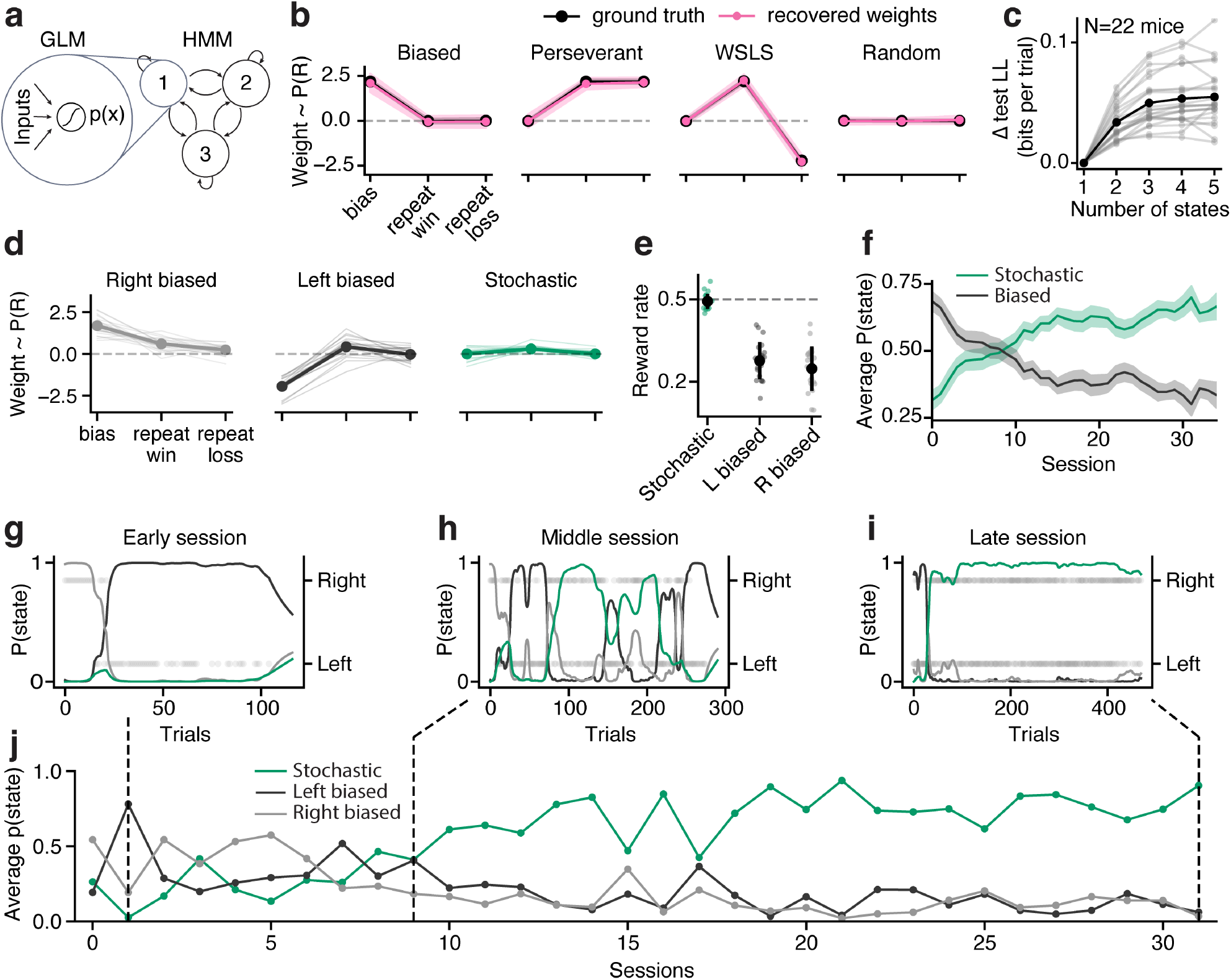
Three-state GLM-HMM describes changes in biased and stochastic strategies across learning. **a**, GLM-HMM (Ashwood et al., 2022) used to identify strategy states across learning. **b**, Recovered GLM weights compared to ground truth for four simulated strategies. **c**, Difference in test log likelihood for GLM-HMM models with one to five states, normalised to the one-state model (n = 22 mice). **d**, GLM weights for a three-state model fit to mouse choices. Thin lines indicate individual mouse fits, thick line indicates mean across animals. **e**, Reward rate of trials belonging to each state where *p*(*state*) > 0.8. **f**, Average P(state) for stochastic and biased (including right biased and left biased) states across learning, averaged across mice. **g-i**, P(state) for stochastic, right biased, and left biased states within example sessions from early (**g**), middle (**h**), and late (**i**) learning. **j**, Average session p(state) of example animal.

We then fit this validated GLM-HMM to mouse data and found that animals’ choices were well captured by a three-state model (Figure 2c-d, Extended Figure 2d), with two states reflecting right-and left-biased strategies. The third state corresponded to a stochastic strategy analogous to the random strategy used in simulation. Trials assigned to the stochastic state achieved a reward rate close to the theoretical optimum of 0.5 (0.49 ± 0.01; n = 22 mice; Wilcoxon signed-rank vs. 0.5; FDR-corrected p = 0.14, Benjamini–Hochberg), indicating that this state captured near-optimal competitive behaviour. In contrast, reward rates of the two biased states were significantly below 0.5 (right-biased, 0.25 ± 0.02; left-biased, 0.28 ± 0.01; Wilcoxon signed-rank vs. 0.5; FDR-corrected p = 7.15 × 10^−7^ for both comparisons; Figure 2e). The probability of occupying the stochastic state increased across learning, consistent with animals progressively adopting stochastic choices as performance improved (Figure 2f-j). Individual session state occupancy revealed a gradual transition across learning stages: early sessions were dominated by the two biased states (Figure 2g), late sessions were predominantly occupied by the stochastic state (Figure 2i), and middle sessions displayed shorter, more interleaved blocks of all three states (Figure 2h). These results suggest that animals did not transition abruptly to optimal behaviour but instead gradually increased use of stochastic strategies while continuing to sample suboptimal biased strategies (Figure 2j). Establishing a framework to estimate trial-by-trial latent states allowed us to go beyond session-based analyses that averaged across heterogeneous behaviours, enabling direct within-animal comparisons of behavioural and neural mechanisms for distinct strategies.

### Comparative analyses of game strategies in monkeys and mice

To further validate our modelling approach and compare game strategies across species, we analysed behavioural data from rhesus monkeys (Lee et al., 2004) playing matching pennies against three computer opponents (Figure 3a-b). The computer algorithms differed in their level of exploitation of the monkey’s past choices and rewards. Algorithm 0 chose both sides with equal probability, such that monkeys could achieve a 50% reward rate regardless of their strategy. Algorithm 1 used 1-4 trial-back choice patterns to predict the monkey’s choice. Algorithm 2 used 1-4 trial patterns of both choice and reward history to predict the monkey’s choice, therefore was able to counter a win-stay/lose-switch (WSLS) strategy. As reported in the original study, monkeys flexibly adapted their strategies against different computer opponents (Figure 3c-e, Extended Figure 3a - h). Monkeys reduced their side biases and the use of WSLS strategy as they progressed through more sophisticated computer opponents that exploited these choice and reward patterns.

**Figure 3.**
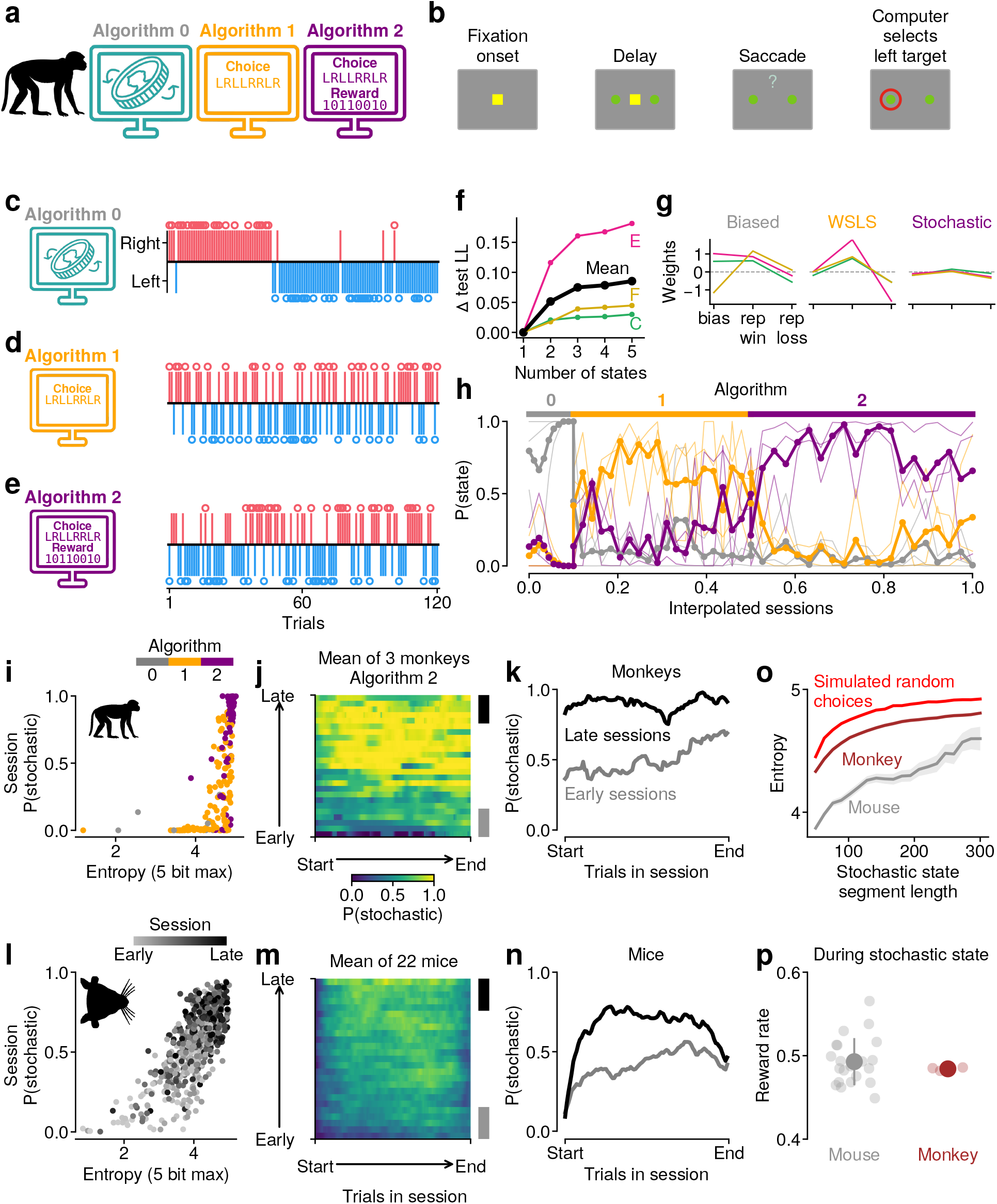
GLM-HMM recovers strategies used by monkeys across algorithms. **a**, Schematic of the three computer algorithms monkeys played against. **b**, Schematic of trial structure in the matching pennies task, adapted from Lee et al., 2004. **c**, Example choices made by a monkey playing against algorithm 0. **d**, Same as c but for algorithm 1. **e**, Same as c but for algorithm 2. **f**, Change in cross-validated log-likelihood as a function of number of states used in GLM-HMM for the monkey dataset. **g**, The weights for the three states obtained. **h**, State occupancy of the three monkeys across the three algorithms. Thin lines denote individual monkeys and thick line denotes mean across monkeys. **i**, Relationship between entropy and P(stochastic) in monkeys across the three algorithms, each dot denotes a single session (logit linear mixed-effects model with random intercept per monkey, p < 0.001, n = 230 sessions from 3 monkeys). **j**, When playing against algorithm 2, the interpolated P(stochastic) in monkeys as a function of the fraction of trials within a session (x-axis) and the fraction of sessions across learning (y-axis). **k**, In monkeys, change in stochastic state occupancy over trials in session in naive (gray) and expert (black). **l**, Same as i but in mice (logit linear mixed-effects model with random intercept per mouse, p < 0.001, n = 639 sessions across 22 mice). **m**, Same as j but in mice **n**, Same as k but in mice. **o**, The estimated 5-bit entropy for different continuous segments of choices within the stochastic state in mice, monkeys and in an agent with simulated random choices. There is a significant difference in entropy across segment lengths between mice and monkeys (0.472 ± 0.17 SE bits, p < 0.001, linear mixed-effects model), and between monkeys and the simulated random agent (0.138 ± 0.0074 SE bits, p < 0.01, linear mixed-effects model). **p**, Reward rate in mice (mean 0.49 ± 0.0059 SEM, n = 22 mice, not significantly different from 0.5, p = 0.238, one-sample t-test) during the stochastic state compared to reward rate in monkeys (mean 0.484 ± 0.0012 SEM, n = 3 monkeys, significantly different from 0.5, p = 0.006, one-sample t-test) during the stochastic state when playing against algorithm 2. There is no significant difference in reward rate between mice and monkeys during the stochastic state (Mann-Whitney U test, p = 0.66, n = 22 mice and n = 3 monkeys).

To formalise the changes in behaviour into single-trial estimates of discrete strategies, we fit the same GLM-HMM used for mice to the monkeys’ choices across all three algorithms. Similar to mice, we found that monkey behaviours can be described by 3 distinct states. Consistent with the model-free analyses, the use of these discrete strategies matched the adaptive behaviours against the corresponding computer algorithms (Figure 3f-h). Monkeys switched from a biased state to a WSLS state when switching from algorithm 0 to algorithm 1, and further switched to a stochastic state upon facing algorithm 2 (Figure 3h). We compared the model performance at predicting out-of-sample choices of both monkeys and mice to other modelling approaches used in the original study (Lee et al., 2004). Our three-state GLM-HMM outperformed the Q-learning model and the logistic regression model incorporating choice and reward history up to five trials back (Extended Figure 4), suggesting that both mice and monkeys’ behaviours may be better characterised by abrupt transitions between distinct strategies than by gradual updating of choice values. Moreover, the logistic regression model revealed only minimal effects of trial history in the stochastic state for both mice and monkeys, despite including regressors five trials into the past (Extended Figure 4g), further justifying the use of 1-trial back regressors in our GLM-HMM.

Inspired by the observation that monkeys flexibly switched their strategies when playing against different computer opponents, we tested whether expert mice that learned to generate stochastic choices could still adapt to changes in competitive pressure. We exposed expert mice trained against Algorithm 2 to interleaved Algorithm 2 and Algorithm 0 sessions (N=4 mice; Extended Figure 5). Because Algorithm 0 does not exploit animals’ choice and reward patterns and chooses left and right randomly, animals could adopt any choice patterns to achieve a 50% reward rate. Remarkably, expert mice with a high level of behavioural stochasticity detected the decrease in competitive pressure within a few trials of the first Algorithm 0 session and adapted accordingly (Extended Figure 5a). Consistent with monkeys’ behaviours, mice only employed the stochastic strategy when needed, and predictable choice patterns significantly increased when simpler strategies such as a strong side bias could achieve comparable reward rates (Extended Figure 5b-f). Although mice rapidly reduced their use of stochastic choices when first exposed to Algorithm 0, repeated Algorithm 0 sessions did not produce a further increase in biased behaviour (Extended Figure 5g-h). Together, we validated the GLM-HMM in identifying dynamic changes in strategies across both mice and monkeys, and demonstrated that mice and monkeys flexibly and selectively recruited stochastic strategies when unpredictability grants a behavioural advantage.

### Mice and monkeys differ in how they deploy the stochastic strategy

The GLM-HMM captured a shared stochastic state between mice and monkeys against Algorithm 2, based on fitted GLM weights. To probe whether this identified state corresponds to the same strategy across species, we examined other model-free behavioural metrics for the stochastic state. In both species, the stochastic state was uniquely characterised by low side bias, high behavioural entropy, and chance-level use of the WSLS strategy (Extended Figure 3i, k). Consistent with this, the stochastic state occupancy per session was positively correlated with choice entropy (Figure 3i, l). To further quantify whether any patterns of the joint animal-computer choice history could predict animal’s next choice, we calculated the mutual information (Methods), and found that the mutual information decreased as the stochastic state occupancy increased (Extended Figure 3j, l). These model-free metrics together support the stochastic state as a shared strategy used by mice and monkeys against an exploitative computer opponent.

We then asked whether monkeys and mice differ in their extent of utilisation of the stochastic strategy and in their extent of “randomness”. Both monkeys and mice increased their stochastic state occupancy across learning when playing against Algorithm 2 (Figure 3j, **m**, Extended Figure 6a, b), but they have different within-session dynamics even during the late stage of learning. Mice were less likely to use the stochastic strategy early during a session, perhaps reflecting a default choice pattern before the computer gains predictive power to pressure the use of a stochastic strategy. Mice were also less stochastic towards the end of a session, reflecting satiation of water reward and potential disengagement (Figure 3n). In contrast, monkeys maintained the use of stochastic strategies throughout each expert session (Figure 3k). Finally, we asked whether mice and monkeys differ in their level of randomness within the respective stochastic state. To this end, we obtained segments of choices where animals were continuously in the stochastic state, and for each segment, computed their 5-bit entropy. To take account of biased estimation of entropy given finite number of trials (Miller, 1955), we computed the 5-bit entropy of simulated random choices to provide a ceiling of the entropy values for each segment length. We found that, controlling for segment length, monkeys exhibited choices of higher entropy than mice, closer to the theoretical maximum (Figure 3o). However, the overall reward rate during stochastic states was very similar across the two species (Figure 3p), suggesting that mice use a lower-entropy regime to achieve comparable reward rates to monkeys.

### Dorsal cortical dynamics encode task-relevant variables and latent strategies

To understand how distributed cortical circuits contribute to structured versus stochastic choices, we performed widefield calcium imaging in transgenic mice expressing GCaMP6s in excitatory neurons while they learned the task (N=7 mice, 28.71 ± 3.45 sessions per subject, 268.21 ± 78.05 trials per session). Widefield signals were corrected for hemodynamic and movement artefacts and registered to the Allen Reference Atlas (Couto et al., 2021), placing all recordings into a common anatomical coordinate space for comparison across mice and sessions (Figure 4a). We then used localised non-negative matrix factorisation (locaNMF) (Saxena et al., 2020) to decompose signals into spatially localised components shared across sessions (Figure 4a-b, see Methods). This approach allowed us to reduce the dimensionality of the widefield signals while preserving spatially meaningful representations of activity across the dorsal cortex. We compared variance explained using locaNMF components versus pixel-averaged atlas brain regions, and verified that locaNMF-based analyses achieved better performance for explaining signal variance and for detecting task-relevant information (Extended Figure 7).

**Figure 4.**
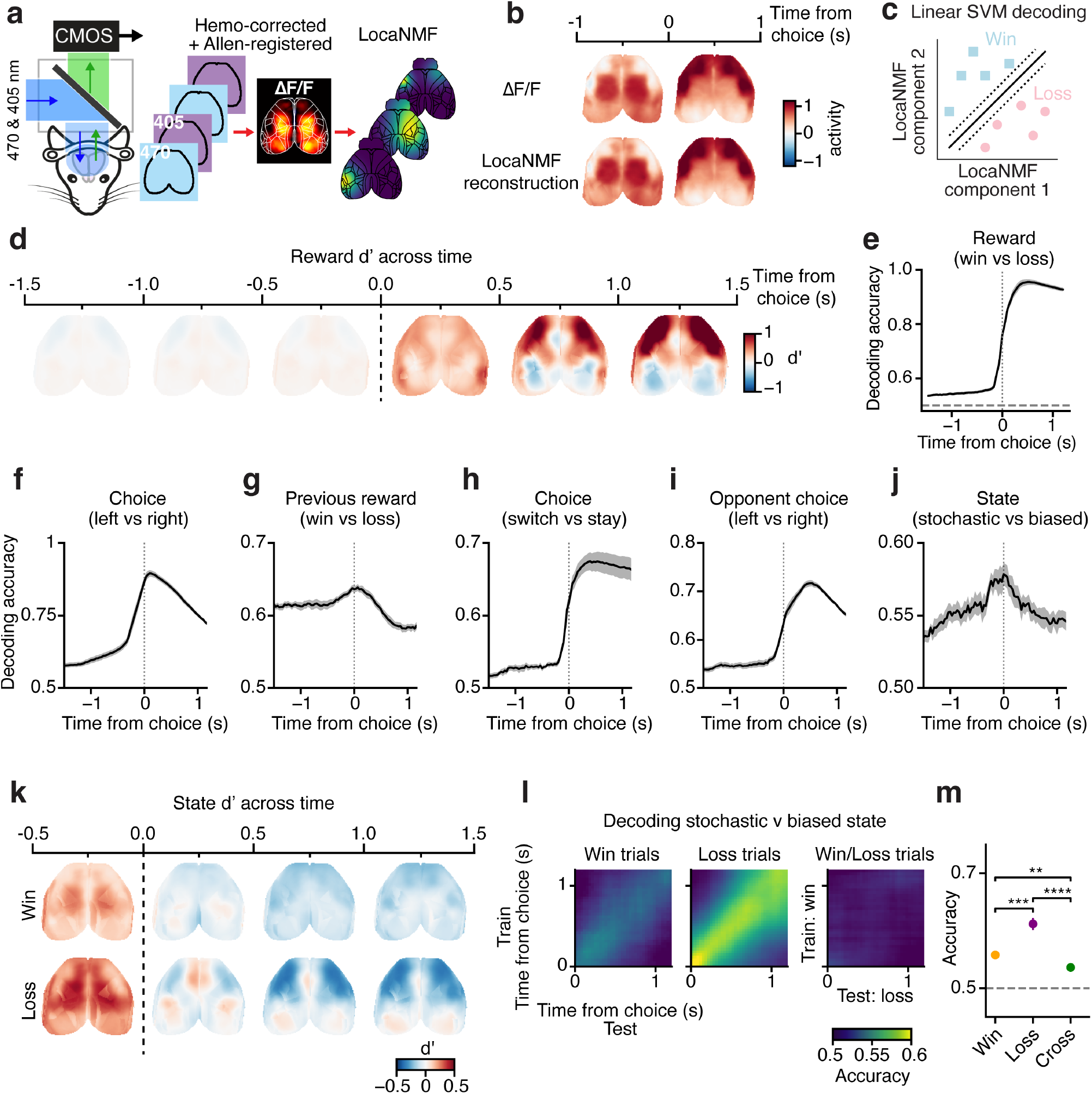
Cortex-wide activity encodes choice, reward, and behavioural state, but the state code is dependent on trial outcome a, Schematic of the widefield acquisition and preprocessing. **b**, Comparison of the ΔF/F widefield signal and the reconstructed locaNMF from an example animal. **c**, Schematic of the use of locaNMF components to perform decoding of task variables using linear SVMs. **d**, d’ of the locaNMF components between rewarded and unrewarded trials. Showing the mean values across 0.5 s time epochs aligned to choice. **e**, Decoding accuracy of reward (win/loss) aligned to choice. **f**, Decoding accuracy of choice (left/right). **g**, Decoding accuracy of previous reward (win/loss). **h**, Decoding accuracy of choice (switch/stay). **i**, Decoding accuracy of opponent choice (left/right). **j**, Decoding accuracy of GLM-HMM state (stochastic/biased). **k**, d’ of GLM-HMM-derived state (stochastic/biased) in rewarded and unrewarded trials. **l**, Cross-decoding analysis showing the accuracy of state decoders trained and tested at different combinations of time windows aligned to choice. Cross-decoding was performed within win trials (left), within loss trials (middle), and trained on win trials and tested on loss trials (right). Accuracy values are the mean across mice per time bin. **m**, Mean peak decoding accuracy of state during the post-choice period (0 - 1 seconds after choice) across mice within win trials, within loss trials, and across win and loss conditions (N=7 mice, paired t-test). Error bars show ± 1 SEM. ** p < 0.01, n.s. not significant.

Using locaNMF components, we related cortex-wide dynamics to task variables using both discriminability index (d’) and linear support vector machine decoding analysis (Figure 4c, see Methods). Dorsal cortical activity contained the key signals necessary to perform the task, including representations of current reward (Figure 4d, e, Extended Figure 7i), spatial choice (Figure 4f, Extended Figure 7j), previous reward (Figure 4g, Extended Figure 7k), switch/stay choice (Figure 4h, Extended Figure 7m), and the opponent’s choice (Figure 4i, Extended Figure 7n). Using a linear encoding model, we showed that these signals were present when accounting for uninstructed movements by the animal, suggesting that these task-relevant representations could not be explained by correlated movement alone (Extended Figure 8). Together, these results demonstrate that distributed dorsal cortical networks encoded information required for both evaluating recent outcomes and generating future choices during competitive gameplay.

Beyond observable task variables, we tested whether cortical activity reflected latent behavioural strategies inferred by GLM-HMM. Early sessions contained more trials in the biased state and late sessions contained more trials in the stochastic state (Figure 2f). To isolate state-dependent signals from time-related fluctuations of neural activity, we sampled equal numbers of trials for the stochastic and biased states within each session before concatenating trials for analyses (Methods). With this balanced dataset, we found robust and dynamic representation of latent behavioural states on the dorsal cortex (Figure 4j,k, Extended Figure 7l). The stochastic state was associated with higher activity than the biased state before choice, but lower activity after choice and during the outcome period (Figure 4k). This strategy-related selectivity was present within each reward condition, but was encoded by distinct patterns of cortical activity across win and loss conditions during the outcome period (Figure 4k-m). Cross-condition decoding revealed that state representations trained on the rewarded trials did not generalise to unrewarded trials (Figure 4l,m). Therefore, cortical circuits appeared to encode latent state in conjunction with reward feedback, consistent with an entangled representation of behavioural strategies and reward contexts.

### Reward history representation is attenuated during stochastic choices

Having established that dorsal cortex encodes reward and behavioural state, we next asked how reward is differentially encoded across these states to support their distinct choice strategies. We formulated two hypotheses for the neural basis of behavioural stochasticity against the exploitative computer opponent, which make opposite neural predictions. In the first hypothesis (H1), mice integrate a longer sequence of choice and reward history than the 1 - 4 trial back n-gram used by the computer in order to effectively counter-predict the computer’s choice (Figure 5a). This requires a strong and persistent memory of trial history and in turn predicts an increase of previous reward information. In the second hypothesis (H2), mice learn to dissociate choice from reward feedback in order to reduce any reward-seeking behavioural patterns that can potentially be exploited by the computer (Figure 5b). This predicts a decrease in previous reward information. To distinguish these hypotheses, we performed decoding analyses of previous reward outcome in the biased and stochastic states, and found a reduction in previous reward information during the stochastic state (Figure 5c - e), consistent with H2. Notably, cross-condition decoding indicates that the form of the previous reward representation is different across states (Figure 5f), suggesting a re-organisation of the neural code in addition to a reduction of selectivity. Further in support of H2, switch versus stay choice information was also reduced in the stochastic state (Extended Figure 9a-c).

**Figure 5.**
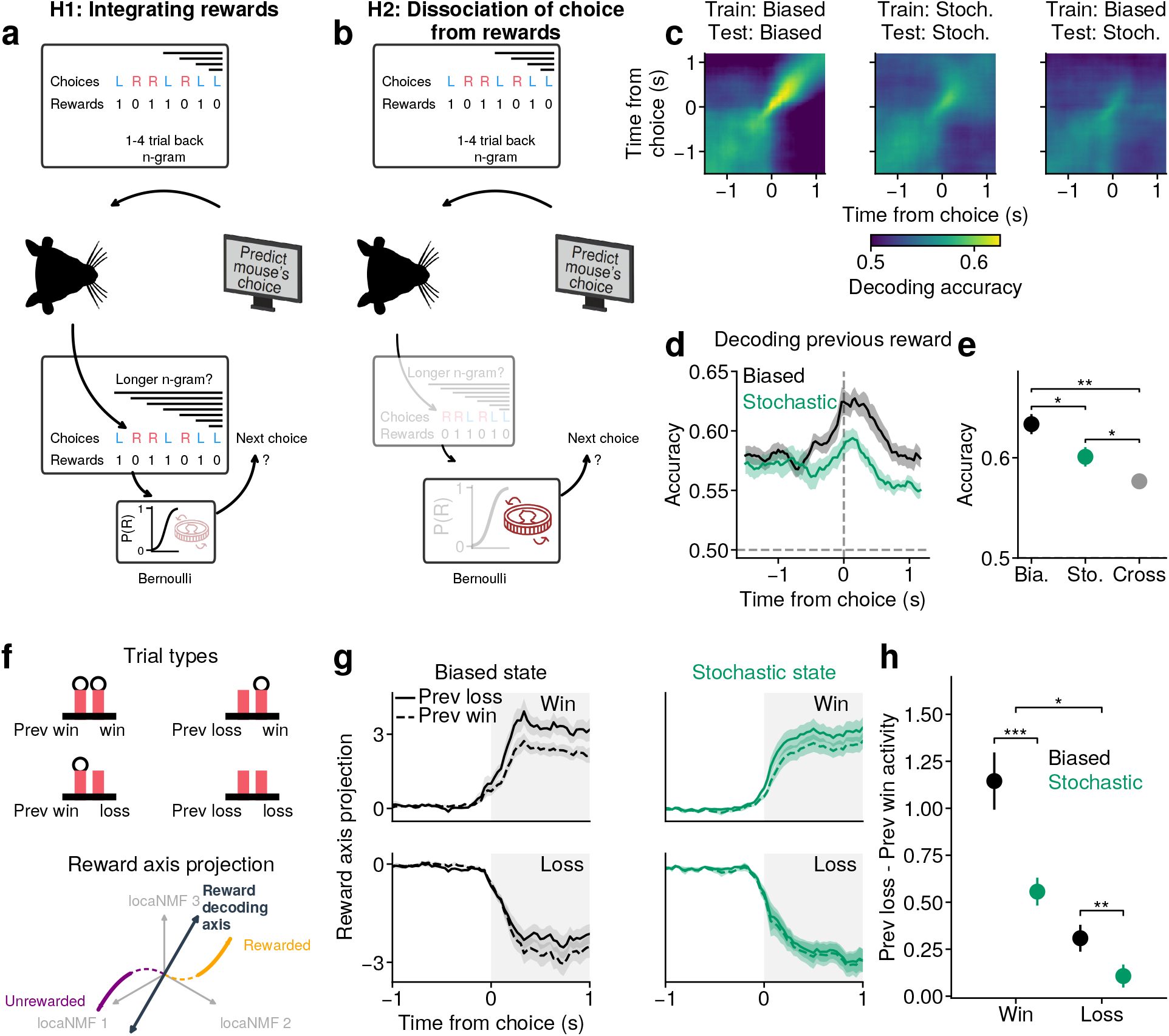
Cortical representation of previous reward is attenuated in the stochastic relative to the biased state. **a**, Hypothesis I of how mice may integrate previous reward and choice information to perform the task by using n-grams longer than the 1 - 4 trial back n-gram used by the computer opponent. **b**, Hypothesis II of how mice may dissociate between previous reward and previous choice with the next choice in order to generate stochastic behaviour. **c**, Cross-window decoding of previous reward within biased states (first column), within stochastic states (second column) and from biased to stochastic states (third column). **d**, Decoding of previous reward during biased and stochastic states. **e**, Summary of the peak of previous reward decoding accuracy across time for within-state and cross-state conditions (paired t-test, n = 7 mice). **f**, Top: schematic of the four possible combinations of previous reward and current reward trials. Bottom: Schematic of the trajectory of neural response projected in relation to the reward decoding axis, dashed part of the trajectories show the activity before choice, and the highlighted part of the line show the activity after choice. **g**, Projection of activity onto the current reward decoding axis, for trials where previous outcome was loss (solid lines) and previous outcome was win (dashed lines), plotted separately for current reward conditions (top row: win, bottom row: loss) and GLM-HMM state (first column: biased states, second column: stochastic state). **h**, Summary of the comparison of previous loss and previous win modulation of current reward activity during biased and stochastic states, using the mean activity in the time period highlighted by the shading in panel **g** (paired t-test, n = 7 mice). * p < 0.05, ** p < 0.01, *** p < 0.001, n.s. not significant.

Beyond decreased history information, a dissociation from reward feedback in the stochastic state also predicts that the response to the current outcome should not depend on what happened on the last trial. To test this, we identified the current reward decoding axis and projected activity from trials where the previous reward outcome was either win or loss (Figure 5f), and compared previous reward modulation of current reward across behavioural states. In the biased state, we found that a previous loss resulted in an amplification of the current win signal, whilst a previous win resulted in an amplification of the current loss signal (Figure 5g, left). In contrast, in the stochastic state, this modulation was significantly reduced (Figure 5g, h). Similarly, while current win signals were stronger following a switch choice than a stay choice in the biased state, this modulation also decreased in the stochastic state (Extended Figure 9d-f). Together, these results suggest that in addition to decreased trial history in the stochastic state, the modulation of current reward outcome by trial history is also reduced to promote stochastic choices.

### Cortical reward signal is decoupled from upcoming choice in the stochastic state

To directly test H2, we asked whether the trial-by-trial variation in cortical reward signals had less predictive power for the upcoming choice in the stochastic state compared to reward-guided states. As a baseline, we isolated choice sequences within the biased state that represented clear reward-guided strategies, which occur when mice switched from one biased state to another. We termed these “block switch” trials (Figure 6a) and verified that mice were indeed more likely to switch choices given a loss and repeat the same choice after a win in these trials (Figure 6b). Decoding analysis confirmed that there was stronger representation of reward history and reward expectation in these block switch trials compared to the stochastic state (Figure 6c,d), whilst current reward decoding remained strong in both states (Figure 6d). We then leveraged single-trial analyses to probe the trial-by-trial relationship between current reward information and the probability of switching on the next trial. Specifically, we trained a decoder to distinguish rewarded and unrewarded trials and projected outcome activity onto the reward decoding axis to provide a read-out of the direction and strength of reward signal on each trial (Figure 6e). We then quantified the correlation between the projected signals on unrewarded trials and the probability of switching on the next trial, separately during block switch, biased, and stochastic states (Figure 6f). As predicted, the reward-guided block switch states displayed the strongest correlation between reward information and the probability of switching, with projections more similar to unrewarded trials leading to higher probability of switching choice in the next trial. This relationship was abolished in the stochastic state (Figure 6g), supporting decoupling between reward and upcoming choice as a mechanism driving stochastic behaviour.

**Figure 6.**
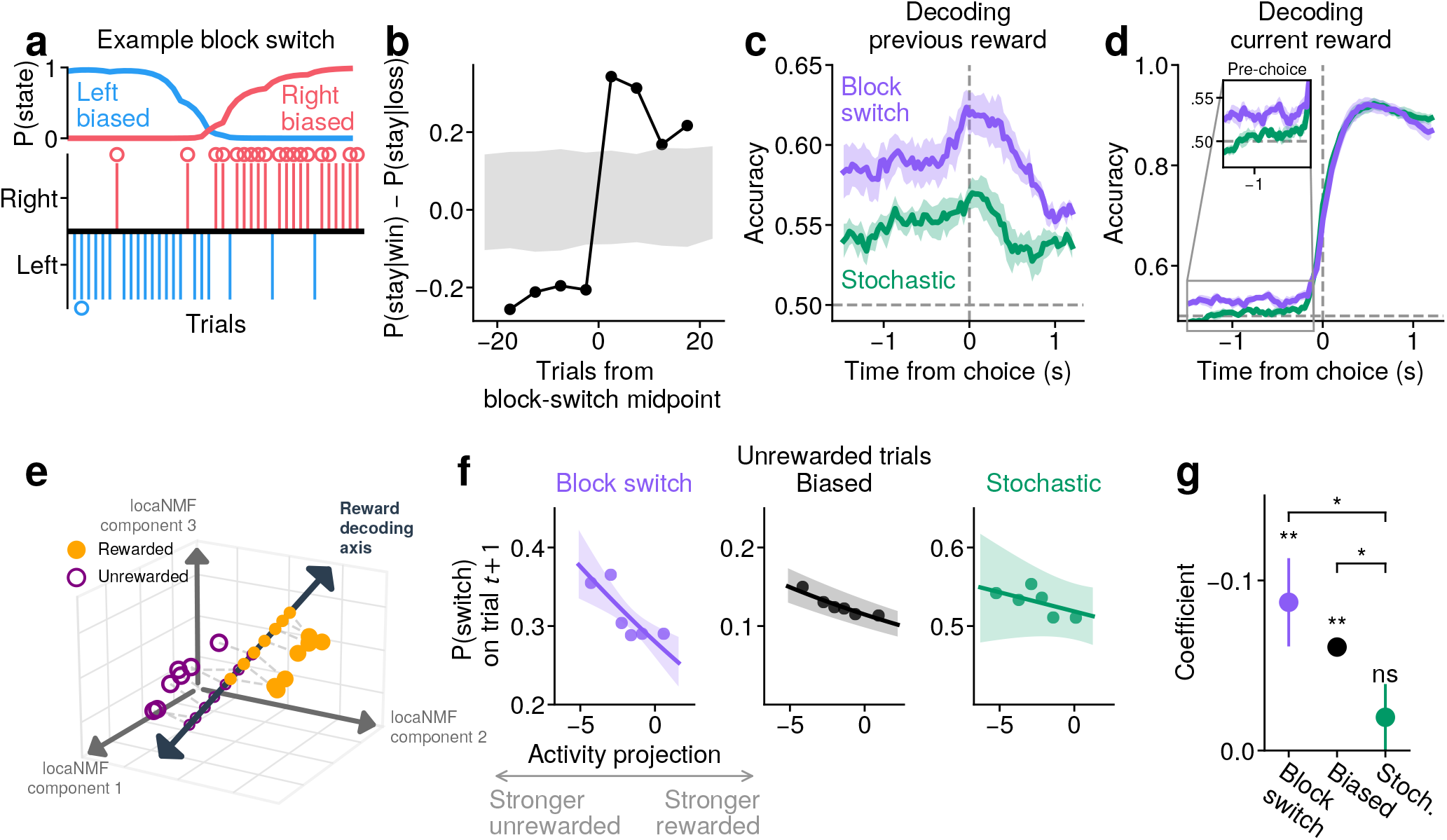
The cortical reward signal is read out to guide choice in reward-sensitive states and decoupled from choice during stochastic behaviour. **a**, Example block switch trial, where an animal shifts from one biased state (left-biased) to another biased state (right-biased). **b**, The difference in P(stay) given a previous rewarded and unrewarded trial aligned to the midpoint of identified block switch trials. Mice are more likely to stay after a loss before the block switch and more likely to stay after a win after the block switch. Shaded gray region shows the 2.5% to 97.5% percentile of values from the null distribution. **c**, Decoding of previous reward information during block switch and stochastic trials. Previous reward decoding was higher during block switch than stochastic trials in both the pre-choice (t < 0 s: 0.591 ± 0.013 vs 0.550 ± 0.007; paired t-test, p = 0.006) and post-choice (t ≥ 0 s: 0.588 ± 0.009 vs 0.546 ± 0.008; p = 0.002) epochs (values mean ± s.e.m., n = 7 mice). **d**, Same as c but for decoding current reward. Current reward decoding was marginally higher during block switch than stochastic trials pre-choice (0.537 ± 0.009 vs 0.514 ± 0.005; paired t-test, p = 0.049) but did not differ post-choice (0.877 ± 0.016 vs 0.884 ± 0.012; p = 0.51; n = 7 mice). **e**, Schematic illustrating the projection of single-trial activity (orange dots for rewarded trials, purple dots for unrewarded trials) onto the current reward decoding axis from widefield locaNMF components. **f**, The probability of choice switching on the next trial given the activity projection onto the reward decoding axis during unrewarded trials, for block-switch, biased, and stochastic state. The line shows the fitted prediction from the generalised linear model and the shaded region shows the 95% confidence interval of the model. **g**, Comparison of the effect of activity projection on P(switch) on the next trial during block switch, biased and stochastic state. Statistical annotation on top of each state shows statistical test for difference from zero, and across states show statistical test for an interaction effect between state and activity projection. * p < 0.05, ** p < 0.01, n.s. not significant.

## Discussion

We studied how mice learned to make unpredictable choices when playing a zero-sum game against a competitive computer opponent (Figure 1). Using GLM-HMM, we found that mice shifted from using predictable, biased strategies to a stochastic strategy as they improved gameplay performance (Figure 2). Applying the same analysis to behavioural data in monkeys playing the same game revealed that this strategy is shared across species, although monkeys had higher choice entropy and more persistent use of the stochastic strategy than mice (Figure 3). We tracked dorsal cortical dynamics as mice learned the task, and found distributed representations of key task-relevant variables, including latent state, choice, reward and previous reward (Figure 4, Extended Figure 7). Reward history and choice information was attenuated and re-organised in the stochastic state compared to the biased states (Figure 5, Extended Figure 9). Furthermore, while the single-trial readout of reward representation was predictive of upcoming choice in reward-guided trials, the correlation between loss information and the probability of switching on the next trial was abolished in the stochastic state (Figure 6). Together, our results provide a putative cortical mechanism for generating stochastic choices through decoupling upcoming choice from reward feedback, in order to avoid forming predictable, reward-seeking choice patterns that can be exploited in multi-agent decision contexts.

Previous characterisations of stochastic behaviour in a similar task have mainly applied model-free metrics such as entropy (Tervo et al., 2014), or reinforcement-learning (RL) models (Lee et al., 2004; Wang et al., 2022). Here, we present the first application of a hidden Markov model, combined with a generalized linear model (GLM-HMM), to describe transitions between discrete states in a competitive game. This model predicted out-of-sample choices better than a reinforcement learning (RL) model and a more complex single-state logistic regression model (Extended Figure 4). This approach allowed us to infer the behavioural strategy on each trial, facilitating more detailed dissection of heterogeneous choice patterns and the underlying neural mechanisms beyond session averages. However, unlike RL-based models (Barraclough, Conroy, and Lee, 2004; Soltani, Lee, and Wang, 2006), GLM-HMM does not provide an explicit learning algorithm from which behavioural variability can emerge. Future studies that combine HMMs with RL models (Venditto et al., 2024), HMMs with higher dimension of latent states across learning (Bruijns et al., 2026; Cuturela and Pillow, 2025), and models with gradual parameter updating rather than abrupt state transitions (Roy et al., 2021) may capture more fine-grained behavioural strategies, although at a potential cost of interpretability.

Using GLM-HMM, we identified dominant strategies of mice and monkey players against an exploitative computer opponent. Early in training, mice more frequently employed a biased strategy compared to monkeys, consistent with previous studies showing strong perseveration (Wang et al., 2022) and dependence on the history of own choices in mice (Wang and Kwan, 2023). In the expert phase, both mice and monkeys converged to a stochastic strategy with low side bias and high choice entropy. While both species achieved comparable reward rate during active engagement against a static opponent (Figure 3p; Algorithm 2), monkeys were more consistently stochastic throughout each expert session (Figure 3i-o), and would likely perform better than mice when facing more sophisticated algorithms. Remarkably, both mice and monkeys flexibly adjusted their levels of stochasticity under different levels of competitive pressure (Figure 3c-h, Extended Figure 5), suggesting that behavioural stochasticity is adaptive and cognitively costly (Baddeley, 1998; Towse and Cheshire, 2007), and would only be employed when needed to avoid exploitation.

Previous studies mostly focused on frontal cortex as a hub for mediating mixed strategies in expert multiplayer games (Abe and Lee, 2011; Barraclough, Conroy, and Lee, 2004; Lee and Seo, 2007; Seo and Lee, 2007; Seo et al., 2014; Tervo et al., 2014; Wang et al., 2025), and for general social cognition across species (Gangopadhyay et al., 2021; Lockwood, Apps, and Chang, 2020). Here, we used an unbiased approach and surveyed the entire dorsal cortex using mesoscale calcium imaging through different stages of learning. Leveraging the novel analytical method of state inference, we identified distributed strategy-related information across multiple cortical regions, in addition to reward and choice signals as observed in other studies (Gauld et al., 2026; Musall et al., 2019; Orsolic et al., 2021; Pinto et al., 2019; Steinmetz et al., 2019). While these task-relevant signals were not uniquely localized to specific brain regions, our survey of the dorsal cortex confirmed stronger representation of reward and latent strategy in the frontal network (Kennerley et al., 2006), consistent with their causal contributions as well as neuromodulatory gating of frontal areas in behavioural variability (Aston-Jones and Cohen, 2005; Tervo et al., 2014).

Beyond the presence of choice, reward, and state representations, we found that the stochastic strategy was associated with attenuated reward history and choice information compared to structured, biased strategies. State modulation of neural responses was not global across all task variables. For example, while reward history was attenuated in the stochastic state (Figure 5), immediate reward information remained strong across states (Extended Figure 9g-i); while switch/stay choice information was reduced in the stochastic state (Extended Figure 9a-c), left/right choice information was not (Extended Figure 9j-l). This selective change of task decodability suggests that differences between biased and stochastic states were unlikely to be explained by correlated changes in arousal or uninstructed movement (Extended Figure 8c-f). Critically, the stochastic strategy was also associated with reduced influence of reward feedback on subsequent choice (Figure 6). This result points towards behavioural stochasticity in a competitive setting emerging, not from counter-predictive strategies, but via a dynamic decoupling of reward signal from choice. While reducing predictability by avoiding reward-seeking behaviours may be a prerequisite for stochasticity, how noise is generated in the brain to guide “random” choices remains to be further investigated. It is possible that counter-predictive and other proposed mechanisms may still drive behaviour in more explicit multiplayer contexts, where an opponent’s choices can be directly observed (Abe and Lee, 2011; Haroush and Williams, 2015; Ong, Madlon-Kay, and Platt, 2021).

Much of neuroscience research has focused on studying how animals learn and take advantage of structures in the environment to guide their actions (El-Gaby et al., 2024). However, probing the neural mechanisms underlying flexible deviation from structure is equally important for understanding adaptive behaviours, especially during social interactions. Failures to recruit adaptive stochasticity are linked to rigid, restricted, and perseverative behaviours observed in neurodevelopmental disorders, compulsive disorders and drug addiction (Gillan et al., 2016; Iversen and Lewis, 2021; Lee, 2013). Therefore, dissecting relevant neural circuits and neuromodulatory systems underlying adaptive stochasticity may open new possibilities for more principled understanding of behavioural inflexibility in psychiatric disorders (Lage, Smith, and Lawson, 2024).

## Methods

### Animals

All experiments were performed in accordance with the UK Home Office regulations (Animals Scientific Procedure Act 1986, PPL: PP4459082) following ethical approval by Sainsbury Wellcome Centre Animal Welfare Ethical Review Body. A total of 24 mice (15 female, 9 male) 8-16 weeks old at the start of experiments were used in this study. 22 mice (10 C57BL/6J (from Charles River Laboratories), 5 Dbh-Cre, 7 Camk2-tTA tetO-GCaMP6s) were used to provide behavioural data for the matching pennies task, and 4 C57BL/6J mice were used for behavioural data for the Algorithm 0 task (see below). 7 mice (Camk2-tTA tetO-GCaMP6s) were used for widefield calcium imaging experiments. Experiments were carried out in the dark phase of the 12/12 h light/dark cycle.

### Surgery

Surgical procedures were carried out within aseptic guidelines. All animals underwent headplate implantation surgery. Mice were induced under anaesthesia using isoflurane (4% induction, 1.5% maintenance) and were given a subcutaneous injection of analgesic (Metacam). The scalp was shaved and cleaned using an antimicrobial skin cleanser (Hibiscrub). Mice were secured onto a stereotaxic frame and placed over a heat pad at 37^∘^C. Temperature was monitored using a rectal probe and eye lubricant was used to protect their eyes. The scalp was cut and removed to expose the surface of the skull, which was cleaned and dried using saline. The skin surrounding the skull was fixed down using an adhesive (VetBond) and the headplate was attached to the posterior end of the skull using clear dental cement (MetaBond). Mice were given a minimum of 7 days to recover from surgery, after which they were water restricted before behavioural training began. Daily water intake was determined by performance in the task (∼1 ml/day). Additional water was provided when necessary to maintain at least 85% of pre-restricted body weight.

### Behaviour set-up

#### Hardware

Mice underwent behaviour training in 50×50×50 cm sound-attenuating aluminium boxes. Mice were head-fixed while sat inside an acrylic tube that allowed for free adjustment of posture. Inside the training boxes, a pair of lick ports were positioned in front of the animal and were attached to two servo-motors that allowed controlled movement in four directions (left, right, forwards, and backwards). The motor could also be manually adjusted upward and downwards. Water delivery was controlled by two solenoid valves that were calibrated to release 1 *μ*l of water per opening. Auditory stimuli were delivered by a front-facing speaker directed towards the mouse. Two infrared-sensitive cameras recorded the front- and side-view of the mouse to capture pupil and orofacial movements during behaviour. An infrared LED and white ambient light were used to illuminate the training box.

#### Matching pennies computer opponent

The computer opponent was designed in line with previous matching pennies algorithms (Barraclough, Conroy, and Lee, 2004; Tervo et al., 2014; Wang et al., 2022). The opponent algorithm predicted the animal’s upcoming choice by exploiting statistical regularities in the history of past choices and outcomes. Specifically, the algorithm tracked the occurrences of all choice-only and choice-reward sequences of length 1-4, as well as an overall choice bias, across the full trial history. For each of these patterns, the opponent evaluated the conditional probability of a leftward choice using a binomial test against the null hypothesis of choosing left by chance (P=0.5), providing nine statistical tests in total. If at least one test reached significance (p<0.05), the opponent based its prediction on the feature associated with the strongest statistical evidence (i.e., the smallest p-value). If no test met the significance criterion, the opponent selected left or right with equal probability.

#### Behaviour training

##### Matching pennies

Mice were first habituated to head-fixation for 3-5 days by slowly increasing the head-fixation time while syringe-feeding water. Following this, the animals underwent two shaping phases. In phase one, animals learned that licking the ports resulted in reward; a water reward of 3 *μ*l per lick was delivered from the chosen spout. A closed-loop detection of a lateralised lick bias (≥ 80% across 20 trials) triggered the lick port motor to shift left-right position by 0.5 mm to counteract the bias. In phase two, animals learned to lick both left and right sides; only one side was rewarded until the animal had chosen it for a total of three trials, after which the rewarded side switched. In this phase, the ports were retracted at the start of each trial and extended within reach during the choice period. Following these two training stages, the animals were trained on the full Matching Pennies task; the trial started with a 10 kHz auditory stimulus that indicated the start of the trial. After a variable delay (1-1.5 s, selected from a uniform distribution), the ports extended and the animal could make a choice within a 2 s choice period. If rewarded, water was delivered immediately after choice, and lick ports retracted after a 2 s consumption period. The inter-trial-interval was drawn from a truncated exponential distribution between 2-5 s (behaviour experiments) or 7-15 s (widefield imaging experiments). On each trial, reward was delivered if the computer opponent predicted the animal’s upcoming choice incorrectly.

##### Interleaving Algorithm 0 sessions

After learning the matching pennies task, some mice played against a non-competitive opponent (Algorithm 0 from Barraclough, Conroy, and Lee, 2004) on alternating sessions, with five sessions of matching pennies interleaved with five Algorithm 0 sessions. This opponent did not track trial history but instead made left/right choices randomly with equal probability.

### Monkey dataset

We used the monkey dataset from Lee et al., 2004 to perform behavioural analysis. The monkey dataset contains behaviour from 3 male rhesus monkeys, 230 sessions and 279,836 trials in total. In their study, they used an oculomotor version of the matching pennies task where monkeys had to saccade towards one of two targets within a 1-second window and maintain fixtation for 0.5 seconds. The monkeys played against three computer algorithms. In Algorithm 0, the computer selected the two choices with equal probability regardless of the monkey’s choice or reward history. In Algorithm 1, the computer uses 1-4 trial-back choice history to makes it choice, and in Algorithm 2, the computer uses 1-4 trial back choice and reward history, analogous to the matching pennies computer opponent used for mice in this study. There are two differences in this algorithm compared to the one mice played against. First, after a significant choice pattern is detected, where the animal is going to choose right with conditional probability *p*, the computer chooses right with probability 1 − *p*, whereas the computer opponent mice played against would have chosen right if *p* ≤ 0.5 and left otherwise. Second, the computer algorithm monkeys played against uses the significant pattern with a conditional probability that is maximally different from 0.5, whereas the computer algorithm mice played against uses the most significant (lowest p value) pattern.

### Behaviour analysis

#### Trial inclusion criteria

For the purpose of mouse behavioural and neural data analyses, we define a miss trial as a trial where no lick was registered during the 2 seconds choice window. These trials were excluded from all analyses.

#### Exponential smoothing function

To approximate reward rate, P(left), and P(Preferred side) within a session, we used an exponential smoothing kernel:

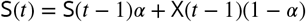

where S is the smoothed result, X is the data (reward or choice), t is the current trial, and *α* controls the weight of the previous trials on the current trial value. We use *α* = 0.95.

#### Entropy

To calculate how equally animals use all three-choice patterns, we used Shannon’s entropy

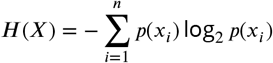

where *p*(*x_i_*) equals the probability of each three-choice pattern, *i*. For 5-bit entropy, we used the five-choice pattern that consists of the animal’s most current choice, previous two choices, and the opponent’s two previous choices.

#### Mutual information

To quantify how much information the joint choice history of the two players contained about the animal’s next choice, we calculated the mutual information between the choices of both players on the two preceding trials and the animal’s current choice. The mutual information is defined as:

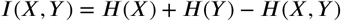

where *X* = (*c_t_*_−1_, *o_t_*_−1_, *c_t_*_−2_, *o_t_*_−2_) are the previous choices of animal *c* and computer opponent *o*. And *Y* = *c_t_* is the current choice of the animal. Mutual information was calculated in blocks of 300 trials with no overlap.

#### Three-choice pattern ranked transitions

To test how predictable the transitions between all three-choice patterns were, we calculated the frequency of transitioning from each three-choice pattern to every other three-choice pattern. This was calculated using a sliding window across three trials and shifting the window by one trial.

### GLM-HMM

We fit a GLM-HMM (Ashwood et al., 2022) to mouse choices to identify distinct strategy states across learning. The model consisted of a hidden Markov model (HMM) that defined the state transitions by a transition matrix *A*, where *A_jk_* = *P* (*S_t_* = *k*|*S_t_*_−1_ = *j*), and a generalised linear model (GLM) that defined the observable choices within each state. Our model was fit using three regressors: general bias, previous-rewarded-choice (R=1, L=-1), and previous-unrewarded-choice (same coding as previous-rewarded-choice). Model parameters were estimated by log-likelihood maximisation using the expectation-maximisation (EM) algorithm; implementation and fitting procedure followed the methods described in Ashwood et al., 2022.

#### Recovery of simulated strategies

To validate that the GLM-HMM can recover strategies in a multi-strategy agent, we tested parameter recovery of a four-strategy agent using simulated data generated from single-state GLMs with known, fixed GLM weights. *(1) Right-biased strategy:* characterised by a strong bias (P(choose right)=0.9) towards the right choice, independent of past outcomes or choices. *(2) Perseverance strategy:* defined by repeating the immediately preceding choice (P(repeat choice)=0.9). *(3) Win-stay/lose-switch (WSLS):* a heuristic learning rule based on the outcome of the previous trial. The rule described repeating your choice if the previous choice was rewarded, and switching if the previous choice was unrewarded (P(WSLS)=0.9). *(4) Random strategy:* an unbiased strategy where the choice is independent of all previous trials and outcomes (P(choose right)=0.5).

The single-state GLMs provided the ground-truth parameters for each strategy. We then tested whether the GLM-HMM could recover these parameters when choices were generated by a multi-strategy agent that transitioned between the four latent states. We simulated 4000-trial sessions from a four-state HMM, where each hidden state generated choices according to the corresponding single-state GLM. The transition matrix was designed with high self-transition probabilities (0.97) and low transition probabilities between states to generate persistent, distinguishable behavioural states (Extended Figure 2a). The initial hidden state was sampled randomly from the four states, after which choices were generated according to the active state and rewards were sampled randomly. On each subsequent trial, the hidden state was sampled according to the transition probabilities, generating the choice and reward sequences used as input for GLM-HMM fitting.

#### Cross-validated model comparison

Model selection across different numbers of states (1-5) was performed using leave-one-session-out cross-validation (CV) for ten different random initialisations, fit separately for each animal. For each CV fold, we separated the data into a training set (all sessions but one) and a test set (left out session) and fit the GLM-HMM to the training set. The trial normalised log-likelihood of the held-out test session was averaged across the ten initialisations. This was repeated across all folds and the results were averaged across all held-out sessions to produce the final test log-likelihood value. The reported metric for model comparison was the test log-likelihood normalised to the 1-state model, indicating the gain in predictive power from incorporating additional hidden states. To obtain a reliable, single set of state-dependent GLM weights per animal, we used a two-stage clustering approach to align and average the recovered parameters across initialisations and sessions. For a single cross-validation fold, the ten sets of GLM weights (across different initialisations) were clustered using constrained k-means to assign the states recovered by different initialisations to the same cluster. The constraint required that each cluster contain the same number of samples (one set of GLM weights per initialisation). We took the median of the weights within each cluster as the representative GLM weight for that state in the left-out session. The median of the state posterior probabilities from the 10 initialisations was calculated for each trial of the left-out session. We used this median P(state) for all subsequent analyses. To derive the final, average GLM weights across all sessions for the animal, the median weights (one set per state from each cross-validation fold) were clustered again using constrained k-means. The mean of the weights in the final clusters was taken as the final set of representative state-dependent GLM weights for the animal. For all analyses that grouped trials into the output states, we used a threshold of P(state) ≥ 0.8.

To estimate P(state) during Algorithm 0 sessions, we used the GLM-HMM fit to choice data from matching pennies (Algorithm 2) sessions to infer latent state probabilities on Algorithm 0 trials.

For fitting GLM-HMM to monkey data, we performed 2-fold stratified cross-validation with stratification such that there is an equal proportion of sessions from algorithm 0, 1, and 2 across folds. We repeated this procedure 20 times and took the mean test log-likelihood over folds.

### Logistic regression model

As a model comparison with our GLM-HMM approach, we fit a logistic regression model (Extended Figure 4) following that from Lee et al., 2004:

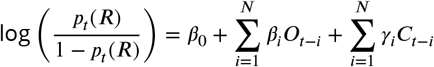

where *p_t_*(*R*) is the probability of choosing right on trial *t*, *β*_0_ is a constant bias term, and *C_t_*_−*i*_ and *O_t_*_−*i*_ are the animal’s own choice and the opponent’s choice *i* trials in the past respectively, each coded +1 for right and −1 for left. *β_i_* and *γ_i_* are the fitted coefficients and *N* = 5 is the number of past trials included. For monkeys, we fit a separate model per algorithm condition (0, 1, 2). For mice, we fit a single model per mouse across multiple sessions. Models were fitted with ℓ_2_-regularised logistic regression with regularisation parameter *C* = 1 and parameters estimated via maximum likelihood.

### Reinforcement learning model

We fit a reinforcement learning model (Extended Figure 4) following that from Lee et al., 2004. On each trial, the model makes choice to the right (*p*(*R*)) or left probabilistically following the difference in value of left and right choices:

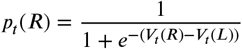

The values are initialised at *V*_1_(*L*) = *V*_1_(*R*) = 0, and updated per trial via

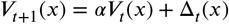

where *x* ∈ [*L*, *R*] is the choice of the animal, *α* is a memory decay term, and the value Δ*_t_*(*x*) for choice *x* is given by:

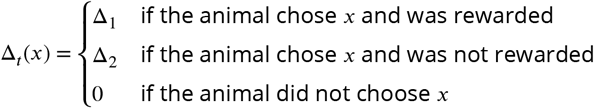

The free parameters (*α*, Δ_1_, Δ_2_) were estimated by maximum likelihood, minimising the negative log-likelihood of observed choices. For optimisation, we initialised the values *α*_0_ = 0.5, Δ_1,0_ = 1.0, Δ_2,0_ = −1.0 and used L-BFGS-B with bounds *α* ∈ (0, 1). Δ_1_ and Δ_2_ values were unconstrained. For model fits to mice data, the parameters were fitted jointly across all sessions for each mouse, with the values *V* resetting back to zero between sessions. For the monkey dataset, we follow the approach in Lee et al., 2004 and fit one set of parameters per monkey per algorithm, and treat all trials as a single continuous sequence, and so did not reset values between sessions.

### Model comparison of GLM-HMM with reinforcement learning and logistic regression models

Since logistic regression models and Q-learning models are not amenable to cross-validation with held-out sessions, as was used in model comparison in Figure 2, we used a within-session cross-validation approach as described in (Ashwood et al., 2022) (Extended Figure 4). For each animal, we performed 5-fold cross-validation across trials to obtain the held-out log-likelihood. The same training and test sets were used for each model. We reported the change in log-likelihood relative to the 1-state GLM-HMM in Extended Figure 4.

### Identification of block switch trials

To identify trial sequences where previous reward information may be used to inform the next choice, we devised a method to identify instances where the animal switches from one biased block (e.g. left-biased) to the opposite biased block (e.g. right-biased), and term these block switch trials (Figure 6). We first identify the biased block using the same P(state) ≥ 0.8 threshold used throughout the study. We then marked the start and end of each continuous sequence of biased states, and identified cases where the beginning of a left-biased block is within 40 trials of the end of a right-biased block (or vice versa). The mid-point of the block switch sequence is then identified as the trial where P(left biased) and P(right biased) are closest. The entire block switch sequence is then defined as 20 trials before and 20 trials after this mid-point. To obtain a null distribution of choice switching behaviour (Figure 6b), we draw the same number of pseudo-midpoints from the same sessions that contributed to the actual midpoints, calculated the same metric over 500 random draws, and computed the 2.5th - 97.5th percentile of those values per bin.

### Defining expert sessions

To compare the stochastic state occupancy and reward rate in mice and monkeys in sessions where they have shown evidence of having learned the task (Extended Figure 6), which we call expert sessions, we first compute the smoothed P(stochastic) across sessions with a centred rolling mean over a 5-session window. We then took the maximum of this smoothed series as an estimate of the P(stochastic) at plateau performance for each mouse. Expert sessions are then defined as sessions where the P(stochastic) is greater than 90% of this maximum and has a value greater than 0.6.

### Widefield imaging

#### Hardware and data acquisition

Widefield calcium imaging was performed using a custom-built tandem-lens epifluorescence macro-scope (85 mm f/1.8D objective, 50 mm f/1.4D tube lens; Nikon), as described previously (Gauld et al., 2026; Orsolic et al., 2021). Two LEDs provided excitation light at 470 nm (M470L4, Thor-labs; excitation filter FF02-447/60-25, Semrock) and 405 nm (M405L4, Thorlabs; excitation filter FF01-405/10-25, Semrock), which were combined via a dichroic mirror (FF458-Di02-25×36, Semrock) and directed onto the sample through a second dichroic (FF495-Di03, Semrock). The two excitation wavelengths were temporally interleaved on alternating frames; the camera acquired frames at 60 Hz, yielding an effective sampling rate of 30 Hz per channel, with LED switching controlled by a Teensy microcontroller synchronised to the camera rolling shutter trigger. Emitted fluorescence was collected through a bandpass emission filter (525/50-25, Semrock) onto a sCMOS camera (pco.edge 5.5, PCO) operating in rolling shutter mode. A custom light-shielding cone was fitted to protect the animals’ eyes from excitation light during imaging. On each trial, imaging was conducted beginning 2 seconds prior to stimulus onset (auditory cue) and continuing for 6 seconds post-stimulus.

#### Data preprocessing

We used the `wfield` package for preprocessing of widefield imaging sessions (Couto et al., 2021). Interleaved 470 and 405 nm frames were separated and concatenated within a session before using SVD to compress the data, keeping only the top 200 components for further preprocessing. Hemodynamic artifacts in the 470 nm channel were corrected by regressing out the 405 nm signal. The corrected 470 nm frames were high-pass filtered before calculating the pixel-wise Δ*F* /*F*_0_, where *F*_0_ was the mean activity across the entire session. Widefield images belonging to the same animal were registered across sessions using a similarity transformation from the ANTsPy package.

All widefield images were registered to the Allen Common Coordinate Framework (CCF) using a landmark-based method to calculate the geometric transformation. A set of corresponding anatomical and functional landmarks were marked on the image space (pixel coordinates of the widefield image) and the atlas space (Allen CCF coordinates). The image space landmarks included visually distinct blood vessel bifurcations (e.g., olfactory bulb intersection with midline) and functionally identified brain regions (e.g., barrel cortex response to whisker stimulation). The corresponding locations were also marked on the Allen CCF reference atlas, forming the set of destination landmarks. The similarity transformation parameters that best mapped the image space coordinates to the Allen space coordinates were determined using a least-squares fit.

### Widefield analyses

#### locaNMF

We performed localised non-negative matrix factorisation (locaNMF) (Saxena et al., 2020) to obtain spatially localised components to perform decoding and regression analyses. Before locaNMF, we performed incremental SVD (Brand, 2002; Brand, 2006) on concatenated widefield data (all sessions per mouse) and took the top 50 SVD components as input to locaNMF. We used a localisation threshold of 90%, a maximum rank per brain region of 3, and a minimum region size of 50 pixels. The number of extracted locaNMF components per mouse ranged from 73 to 80.

#### Comparison of locaNMF with SVD and averaging pixel across brain regions

To validate the locaNMF method, we first compared the signal variance captured by locaNMF compared to performing SVD on the widefield dataset and taking the mean pixel value per atlas-defined brain region. We performed incremental SVD on the session-concatenated widefield data and took the 500 SVD reconstruction as the signal. We then compared the variance explained using increasing number of brain region averages (Extended Figure 7a), SVD components (Extended Figure 7b) and locaNMF components (Extended Figure 7c). We found that 50 SVD components were sufficient to capture over 99% of the signal variance. Using those components as input, locaNMF reached a similar level of explained variance, with considerably lower signal variance explained by taking average pixel values per brain region (Extended Figure 7d).

We also compared the stability of the decoding axis obtained using locaNMF compared to per-region pixel averages. We trained decoders to decode reward in 3 sets of sessions throughout the learning of an example mouse, and found that the reward axes are more spatially consistent across sessions when using locaNMF compared to using per-session pixel averages, reflecting potential variances in recording quality across sessions that are denoised by performing session-concatenated SVD (Extended Figure 7e, g). However, the time courses of the activity projected onto the reward axis are similar (Extended Figure 7f, h), consistent with strong reward representation in the dorsal cortex throughout learning. Across mice, we compared the decoding performance of reward, choice, previous reward, state and switch/stay using locaNMF components and using per-session pixel averages, and found that locaNMF generally gives superior performance (Extended Figure 7i-n).

#### Discriminability (d’) index

Trial-by-trial neural discriminability was assessed on the temporal components produced by lo-caNMF. Each recording session was first z-score normalised. For any two trial conditions A and B, discriminability was quantified with the d’ sensitivity index:

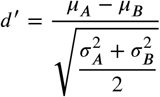

where *μ* and *σ* were the across-trial mean and standard deviation of z-scored component activity for each condition. Prior to computing d’, single-trial component activity was first averaged within each 0.5 s epoch spanning -1.5 to +1.5 s relative to lick onset; d’ was then computed directly on these epoch means. The component-wise d’ was projected into pixel space as the weighted sum of d’ across components (using each component’s spatial weights), producing pixel-wise d’ maps.

Trials in each condition were balanced before computing d’: all trials from the minority condition were retained, and an equal number of trials was randomly subsampled from the majority outcome. d’ was then computed on each size-matched subsample. We repeated subsampling 50 times and averaged across all repeats to obtain the reported d’ estimates. In the case where discriminability was assessed within the stochastic and biased GLM-HMM states, trials were first balanced within each session to match the number of biased and stochastic trials. This prevented individual sessions with a strong state imbalance from disproportionately influencing the results.

#### Comparison across states in widefield analysis

For the purpose of comparison across state in the widefield analysis, we used the same P(state) ≥ 0.8 threshold as in the behavioural analysis to obtain trials of stochastic, left-biased and right-biased state. Left-biased and right-biased state were then combined to form a single biased state to compare against the stochastic state.

#### Widefield decoding

We used the extracted locaNMF components described in the previous section to perform decoding of GLM-HMM state (Figure 4j-m), current reward (Figure 4e), previous reward (Figure 4g), spatial choice (Figure 4f), switch versus stay choice (Figure 4h) and opponent spatial choice (Figure 4i). We first aligned the extracted activity to 1.5 seconds before and after the first lick event. For each 0.25 second time bin within this window (with 0.03 seconds time-steps, constrained by the 30 fps sampling rate of widefield imaging), we took the mean activity across time such that we have a population vector per trial to perform decoding.

For general decoding per task variable (Figure 4e-j, Extended Figure 7i-n), we balanced trials across conditions as follows: for current reward and previous reward decoding, we balanced trials across both of these variables; for choice decoding, we balanced choice and reward; for GLM-HMM state decoding, we balanced state and reward, and subsampled trials from each session such that there was an equal number of stochastic and biased trials per session; for switch versus stay and opponent choice decoding, only trials across the decoded variables were balanced. For decoding of states within each rewarded or unrewarded condition, (Figure 4l, m), we subsampled trials from each session such that within each reward condition, there was an equal number of stochastic-and biased-state trials from each session. For comparison of decoding in different states, we subsampled trials from each session such that there was an equal number trials from each state from each session. Following this state subsampling procedure, for decoding of task variables within and across GLM-HMM states, we performed a 2 by 2 balancing such that there was both an equal number of trials from each state condition and that the task variable classes were balanced within each state condition. Sessions with fewer than 10 trials in any of the conditions were excluded from the analysis. We used an ℓ_2_-regularised linear SVM decoder with regularisation parameter *C* = 1, and reported the mean decoding accuracy using 2-fold cross-validation (and 2-fold cross-validation for decoding involving block switch states, due to reduced trial counts) repeated 20 times.

For cross-condition decoding, (Figure 4l, m, Figure 5c, e, Extended Figure 9b, c, h, i, k, l), states were first split using the P(state) ≥ 0.8 threshold. We then use the same time window and time bin as described but now train a decoder on activity in one time window, and test it on every other time window. For within-condition decoding, we used the same 2-fold cross validation on held-out folds. For across-condition decoding, we tested on all trials of the target condition.

#### Reward activity projection analysis

To investigate the relationship between reward-related activity and probability of switching choices in the next trial (Figure 6e-g), we first trained separate ℓ_2_-regularised linear SVM decoders to decode reward in the stochastic state, biased state and block switch trials in each time window aligned to the time of choice. We then took the dot-product of the lick-aligned locaNMF activity at each time window and the corresponding decoder weight vector. This was done in a cross-validated fashion: each trial’s projection is calculated from the dot-product of weights with a decoder trained on a held-out dataset, and the projection values are averaged across folds. To obtain a single value per trial to relate with the probability of switching in the next trial (Figure 6e), we took the projection values averaged over the post-lick window 0.5 - 1.0 s. We specifically looked at projection values during the unrewarded trials to ensure that differences in probability of switching in the next trial cannot be attributed to the actual reward received, but is due to the difference in reward-related activity. Statistical comparison of the relationship between projection values and P(switch) was performed using a logistic generalised linear model with cluster-robust standard errors grouped by subject. For visualisation in Figure 6f, we plot the fitted logistic curve and the datapoints binned into 6 projection quantiles. For each state, we tested whether the slope coefficient is significantly different from 0, and tested the interaction of state and projection effect to test for significant differences in the effect of the projection value between states.

### Widefield encoding model

To quantify the contributions of task variables and animal movements on widefield activity, we fit encoding models to the widefield data (Extended Figure 8). For each mouse, we fit a ridge regression model (with regularisation parameter *α* = 1) with the following regressors: task events (sound onset, port extension, and port retraction, each with a temporal kernel from 0 to 1 second aligned to the onset of the corresponding event); behaviour (time of first lick and lick direction, each with a temporal kernel of -2 to 2 seconds); movement (z-scored motion energy of the face camera, resampled to match the sampling rate of the widefield recording); reward and previous reward (temporal kernel of -2 to 2 seconds aligned to first lick); and strategy (either stochastic or biased GLM-HMM state, with a temporal kernel of -2 to 2 seconds aligned to first lick). The temporal kernels were constructed by a set of time-shifted binary regressors, one per imaging frame within its temporal window. Since the weights are independent from each other, the temporal kernels were not constrained and can take any shape.

To compare the variance explained by individual features alone (Extended Figure 8b), we fit separate ridge regression models and computed the mean across mice. Variance explained measures were weighted by the variance of each locaNMF component and evaluated -1 to 5 seconds relative to sound onset pooled across trials. To compare the unique variance explained by individual features (Extended Figure 8c), we used a leave-one-out method and computed the variance explained by the full model subtracted by the variance explained by a model with the feature of interest left out. To obtain the lick-aligned variance explained (Extended Figure 8d-f), we re-aligned the model prediction to lick onset and computed the variance explained in 500 ms windows from -1 to 2 seconds relative to lick, pooled across trials and time frames. To obtain the spatial map of variance explained, we multiply the variance explained per component by the spatial weight of each locaNMF component.

## Competing interests

The author declare no competing interests.

## Acknowledgment

This work was funded by the Sainsbury Wellcome Centre Core Grant from the Gatsby Charitable Foundation (GAT4057) and Wellcome (318818/Z/24/Z), by a UK Research and Innovation grant (EP/Y008804/1); and by the Gatsby Initiative in Brain Development and Psychiatry (GAT3955). We thank members of the Duan laboratory and members of the Erlich laboratory for discussion and advice; J. Erlich for help to implement the computer opponent; J.W. Pillow and P. Dayan for discussion on early versions of GLM-HMM; SWC Neurobiological Research Facility for animal support; and FabLabs for contributions to hardware design.

## Author contributions

C.A.D. and J.A. conceived the project. J.A. and C.A.D. designed the behavioural setup and task, with assistance from O.M.G. J.A. performed surgeries. J.A. and M.M. contributed to mouse behavioural training. J.A. collected widefield data. J.A. and T.P.H.S. analysed mouse behavioural and widefield data. D.L. provided the monkey behavioural data. T.P.H.S. analysed monkey data. J.W. implemented the GLM-HMM. C.A.D. provided supervision and input to all data analysis. C.A.D., J.A. and T.P.H.S. wrote the manuscript, with comments from co-authors.

## Data and code availability

Code and data related to this manuscript are available at: https://gin.g-node.org/DELab/matching-pennies-widefield-2026

---

**Extended Figure 1.**
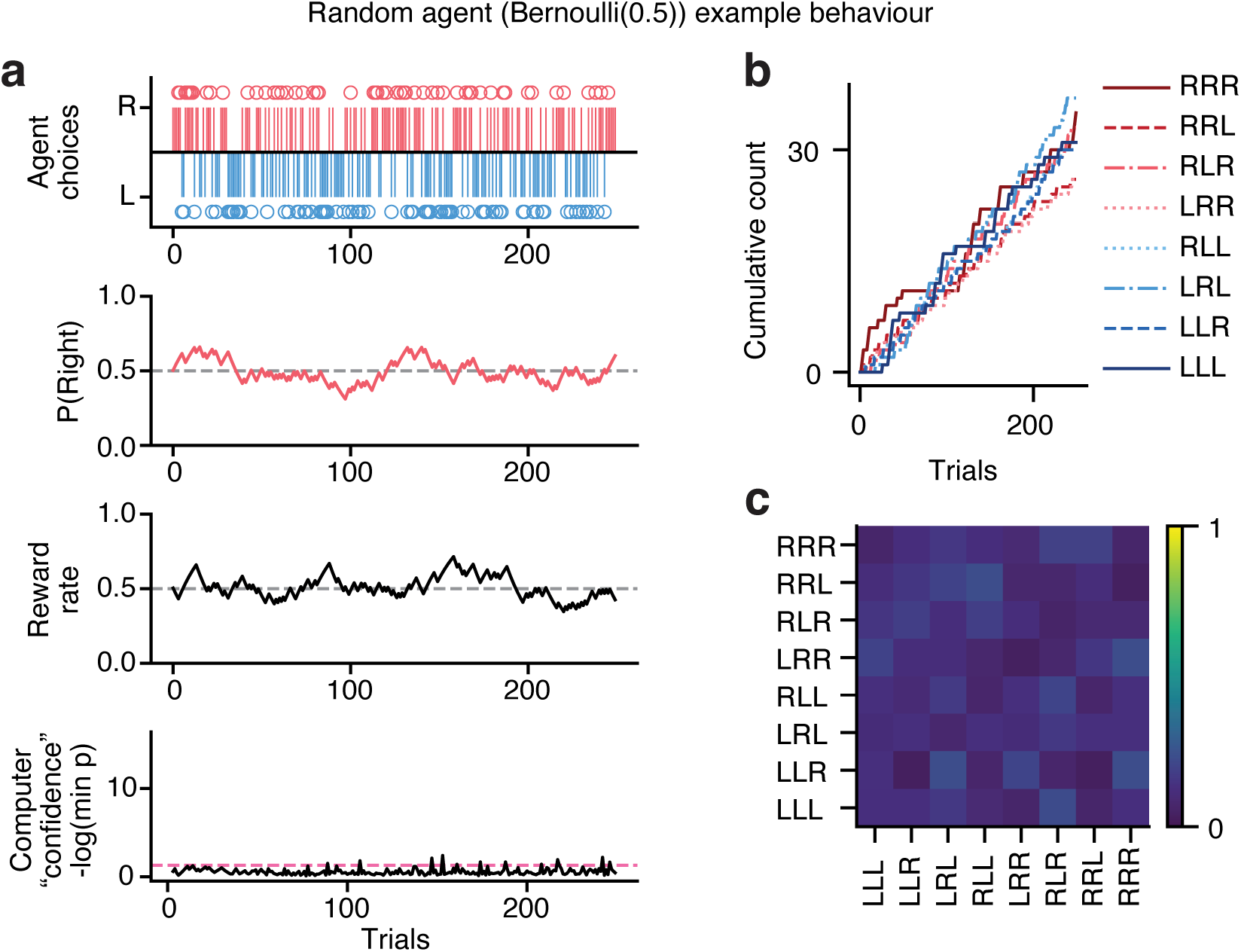
Bernoulli(0.5) simulated agent plays the matching pennies game. **a**, Top row, agent choices, circles indicate rewarded trials. Second row, smoothed probability of right choice. Third row, smoothed reward rate. Bottom row, negative log of the minimum p value. Dashed pink line indicates where p = 0.05. **b**, Cumulative count of all 2^3^ possible three-choice patterns. **c**, Transition matrices showing the ranked frequency of transitions between all 3-choice patterns.

**Extended Figure 2.**
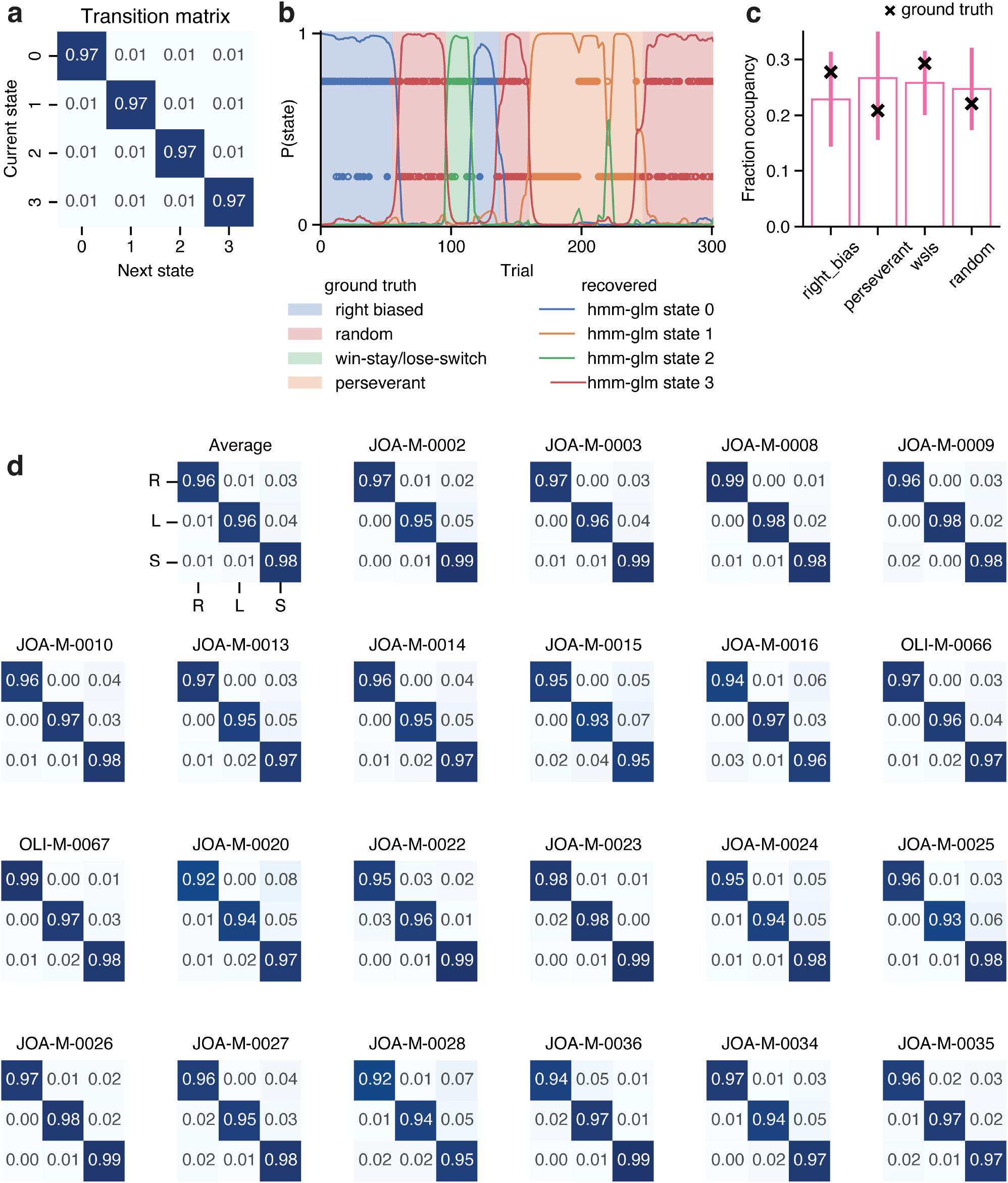
Validation of GLM-HMM using simulated data. **a**, Transition matrix for a four-strategy simulated agent. **b**, Recovered p(state) of GLM-HMM fit to simulated four-state choices (lines) compared against the ground truth (coloured blocks), shown for a subsection of trials. Filled circles indicate rewarded choices, empty circles indicate unrewarded choices. **c**, Recovered fraction occupancy for each state across 10 model fit runs. **d**, Transition matrix for GLM-HMM fit to mouse data.

**Extended Figure 3.**
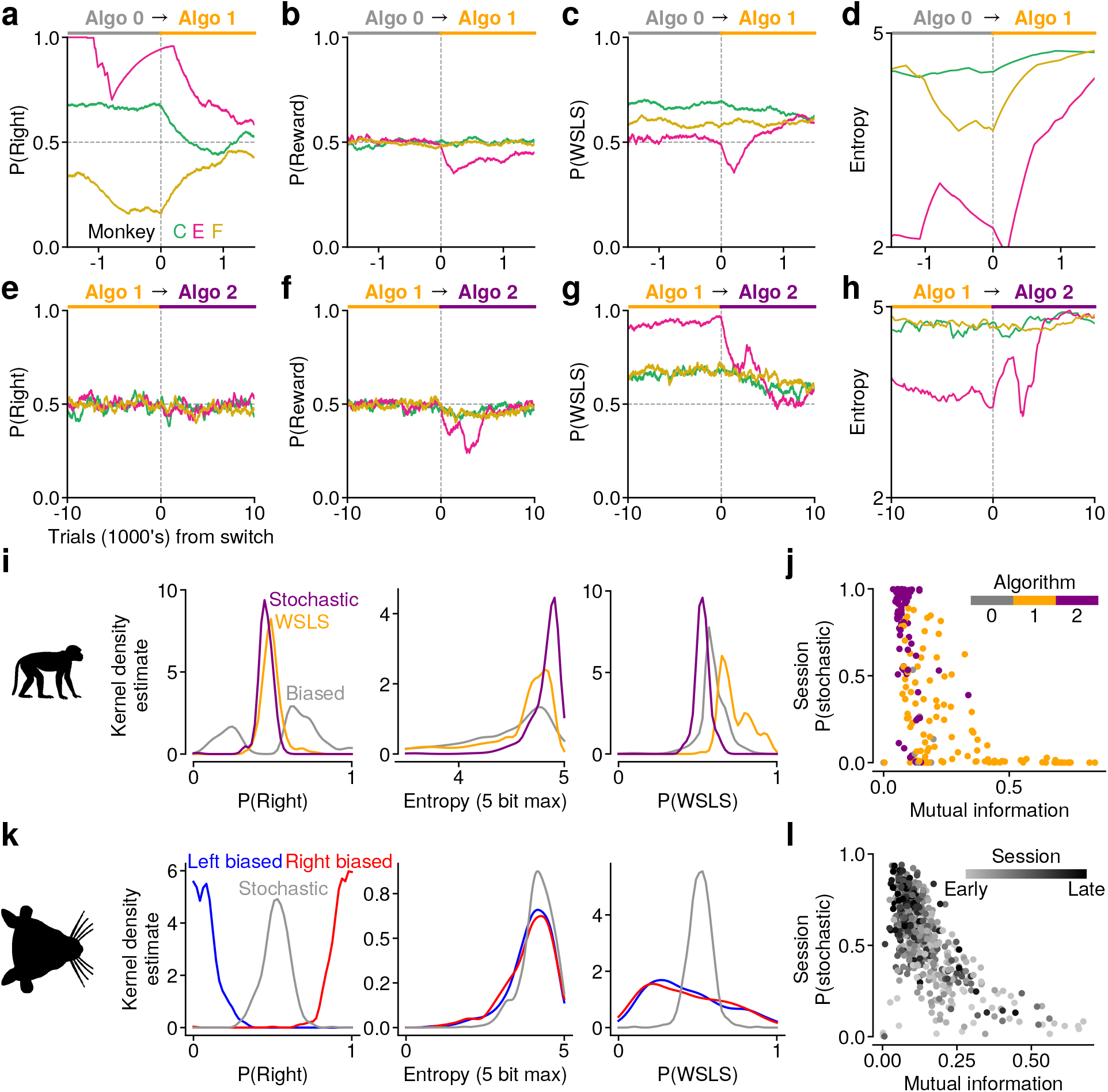
Model-free metrics validate the stochastic state as a genuine signature of unpredictable choice across species. **a**, Change in P(right) from algorithm 0 to algorithm 1. **b**, Same as a but for P(reward). **c**, Same as a but for P(WSLS). **d**, Same as a but for entropy. **e**, Change in P(right) from algorithm 1 to algorithm 2. **f**, Same as e but for P(reward). **g**, Same as e but for P(WSLS). **h**, Same as e but for entropy. **i**, Kernel density estimate of P(right), entropy and P(WSLS) across the three states in monkeys. **j**, Relationship between mutual information and P(stochastic) in monkeys, each dot denotes a single session (Spearman’s *ρ* = −0.71, p < 0.001, n = 230 sessions). **k**, Same as i but in mice. **l**, Same as j but in mice (Spearman’s *ρ* = −0.62, p < 0.001, n = 639 sessions, 22 mice).

**Extended Figure 4.**
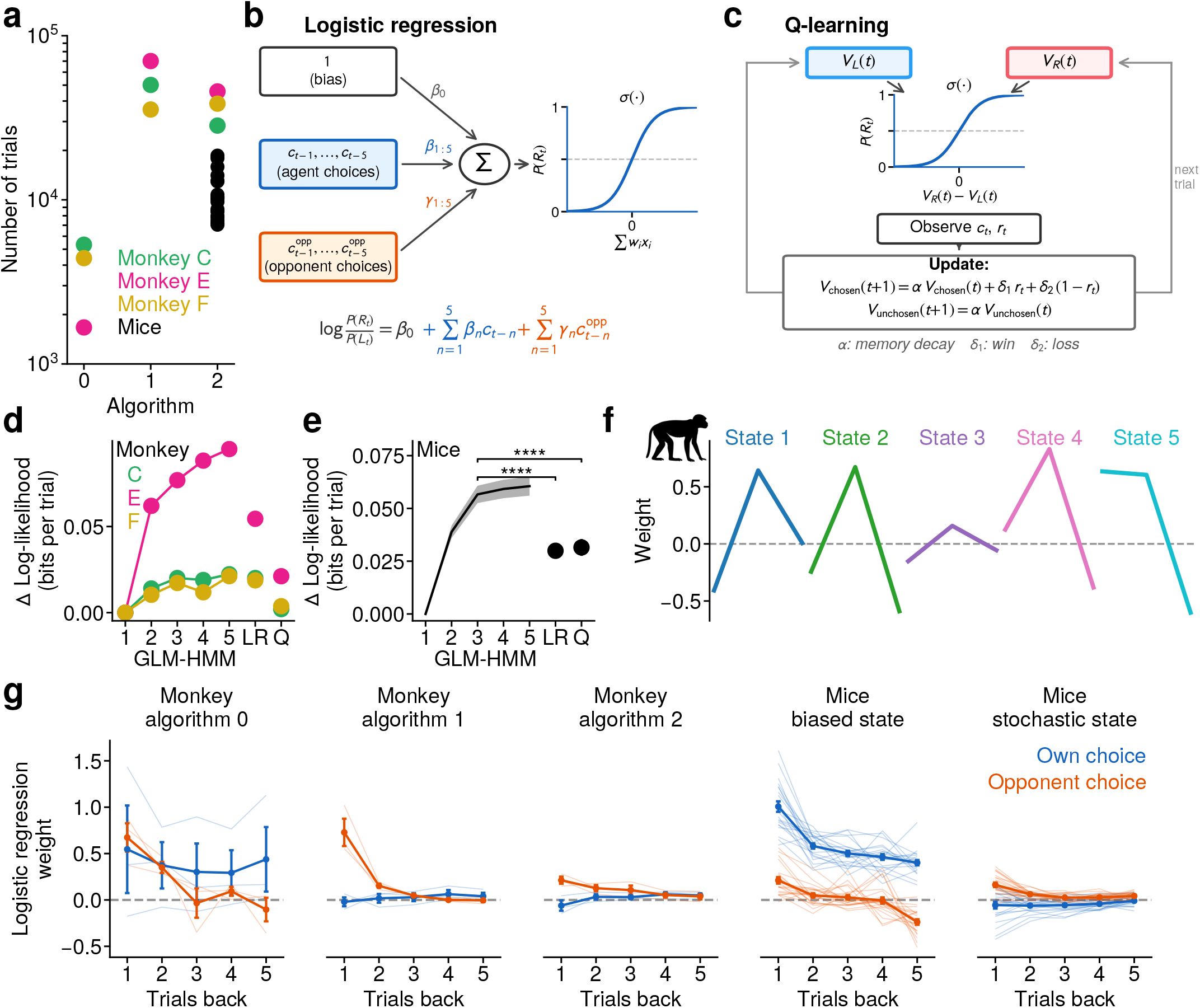
Further comparison of mouse and monkey behaviour. **a**, Number of trials each monkey and mouse was trained on for each computer algorithm (n = 3 monkeys, n = 22 mice). **b**, Schematic of the logistic regression model used in (Lee et al., 2004) to model animal choice. **c**, Schematic of the reinforcement learning model used in (Lee et al., 2004) to model animal choice. **d**, In monkeys, cross-validated log-likelihood comparison between GLM-HMM with different number of states, logistic regression (LR) model and reinforcement learning model (Q). **e**, Same as d but in mice (paired t-test, n = 22 mice). **f**, Weights for each state obtained when fitting a 5-state GLM-HMM to monkey E. **g**, Weights from the logistic regression model fitted on monkey behaviour against different computer algorithms (first three columns), and fitted on mouse behaviour separately within the biased and stochastic states (last two columns; state assigned per trial from the GLM-HMM posterior, with regressors lagged within contiguous same-state blocks; n = 22 mice). * p < 0.05, **** p < 0.0001

**Extended Figure 5.**
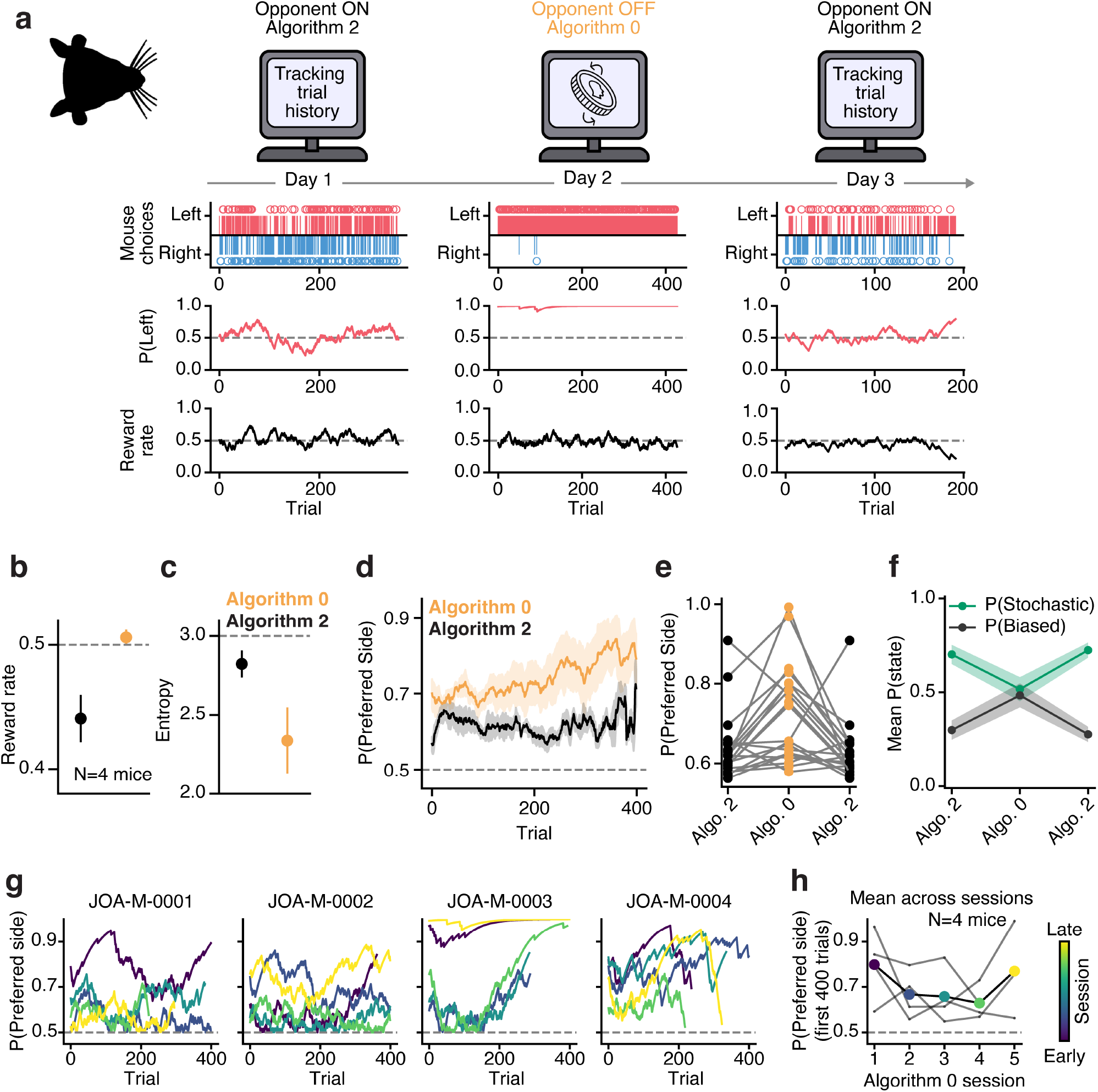
Mice flexibly switch strategies in response to change in competitive pressure. **a**, Example choice behaviour as the mouse alternate playing a predictive computer opponent (Algorithm 2, left and right columns) and a computer opponent choosing both sides with equal probability (Algorithm 0, middle column). **b**, Reward rate of mice playing against Algorithm 2 and Algorithm 0. **c**, Same as b for 3-bit entropy. **d**, Moving window average of the probability of biasing towards any preferred side within sessions comparing play against Algorithm 0 versus Algorithm 2. **e**, Probability of biasing to a preferred side across three consecutive sessions, switching from Algorithm 2 to Algorithm 0 and back to Algorithm 2. Includes data across all sessions and all mice (N=4 mice, 20 triplet sessions) **f**, Mean probability of the GLM-HMM stochastic and biased state across triplet sessions, averaged across all sessions and mice. Welch’s two-sample t-test comparing P(Stochastic) in Algorithm 2 versus Algorithm 0 sessions: t(34.25)=2.48, p=0.018; mean P(Stochastic) against Algorithm 2: 0.705 ± 0.208, n=24 sessions; Algorithm 0: 0.517 ± 0.283, n=20 sessions. **g**, Moving window of the probability of biasing towards a preferred side within sessions when playing against Algorithm 0, coloured by session order. **h**, Probability of biasing towards a preferred side when playing against Algorithm 0, ordered by session order and averaged across mice (N=4 mice).

**Extended Figure 6.**
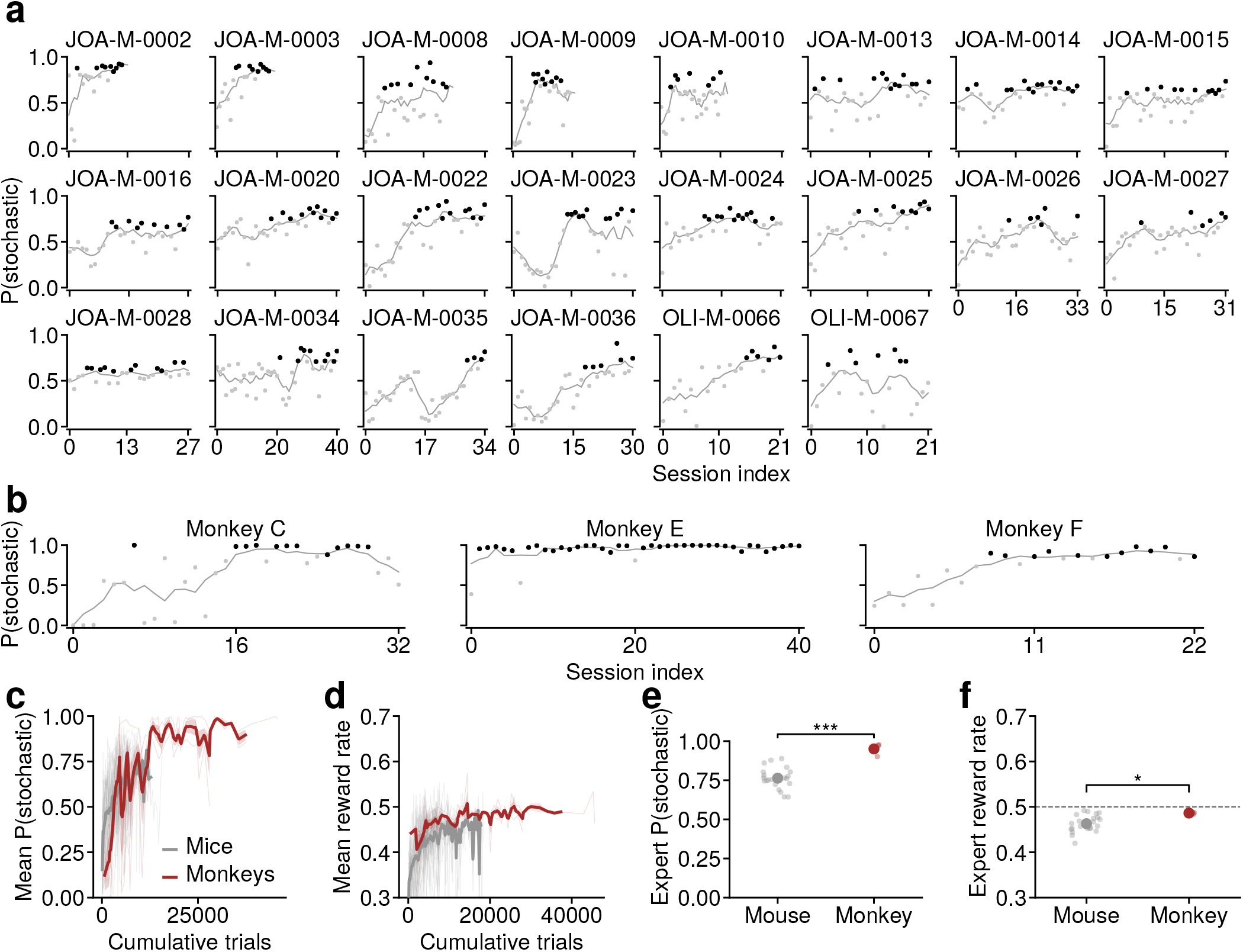
Change in stochastic state occupancy and reward rate in mice and monkeys over learning. **a**, Stochastic state occupancy as a function of chronologically sorted sessions, gray line indicates a rolling mean and dots in black indicate expert sessions (see Methods). **b**, Same as a but in monkeys. **c**, Stochastic state occupancy as a function of cumulative trials computed on each session; thick lines indicate the mean for mice (gray, n = 22) and monkeys (brown, n = 3). **d**, Same as c but for reward rate. **e**, The mean stochastic state occupancy in expert sessions in mice (n = 22, 0.763 ± 0.014) and monkeys (n = 3, 0.950 ± 0.024; p < 0.001, Mann-Whitney U test). **f**, Same as e but for reward rate. There is a significant difference in reward rate between mice (n = 22, 0.463 ± 0.004) and monkeys (n = 3, 0.486 ± 0.001; Mann-Whitney U test, p = 0.036). Dashed line indicates the Nash equilibrium reward rate of 0.5. * p < 0.05, *** p < 0.001

**Extended Figure 7.**
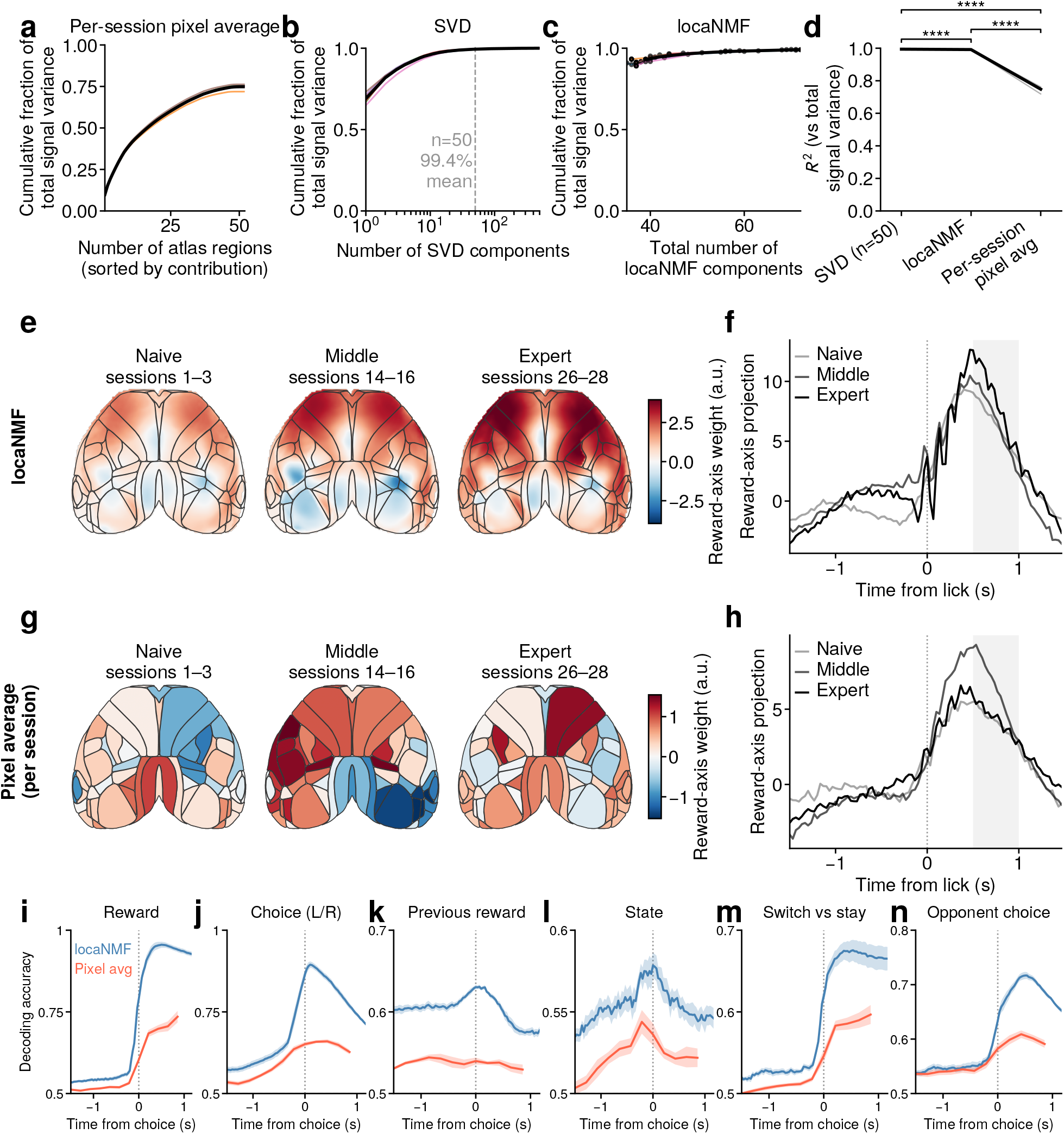
Comparison of locaNMF with per-session brain region averaging. **a**, Cumulative fraction of total signal variance as a function of the number of atlas regions used via averaging widefield signals per region per session. **b**, Same as a, but as a function of the number of SVD components used. **c**, Same as a, but as a function of the number of locaNMF components using the 50 SVD components as input. **d**, Comparison of the percentage of total signal variance explained using SVD with 50 components, locaNMF, and per-session pixel averaging of atlas regions (n = 7 mice). **e**, Weights for reward decoding using locaNMF signals and an SVM decoder, obtained in 3 sets of sessions for an example mouse. **f**, Projection of weighted activity during rewarded trials during the three sets of session shown in e. **g**, Same as **e**, but averaging widefield signals across pixels per brain region per session. **h**, Same as f, but averaging widefield signals across pixels per brain region per session. **i**, Comparison of reward decoding accuracy across all trials concatenated across sessions using locaNMF components and using per-session pixel averages across brain regions. **j**, Same as i but for choice decoding. **k**, Same as i but for previous reward decoding. **l**, Same as i but for decoding of GLM-HMM state (stochastic versus biased). **m**, Same as i but for switch versus stay decoding **n**, Same as i but for opponent choice decoding.

**Extended Figure 8.**
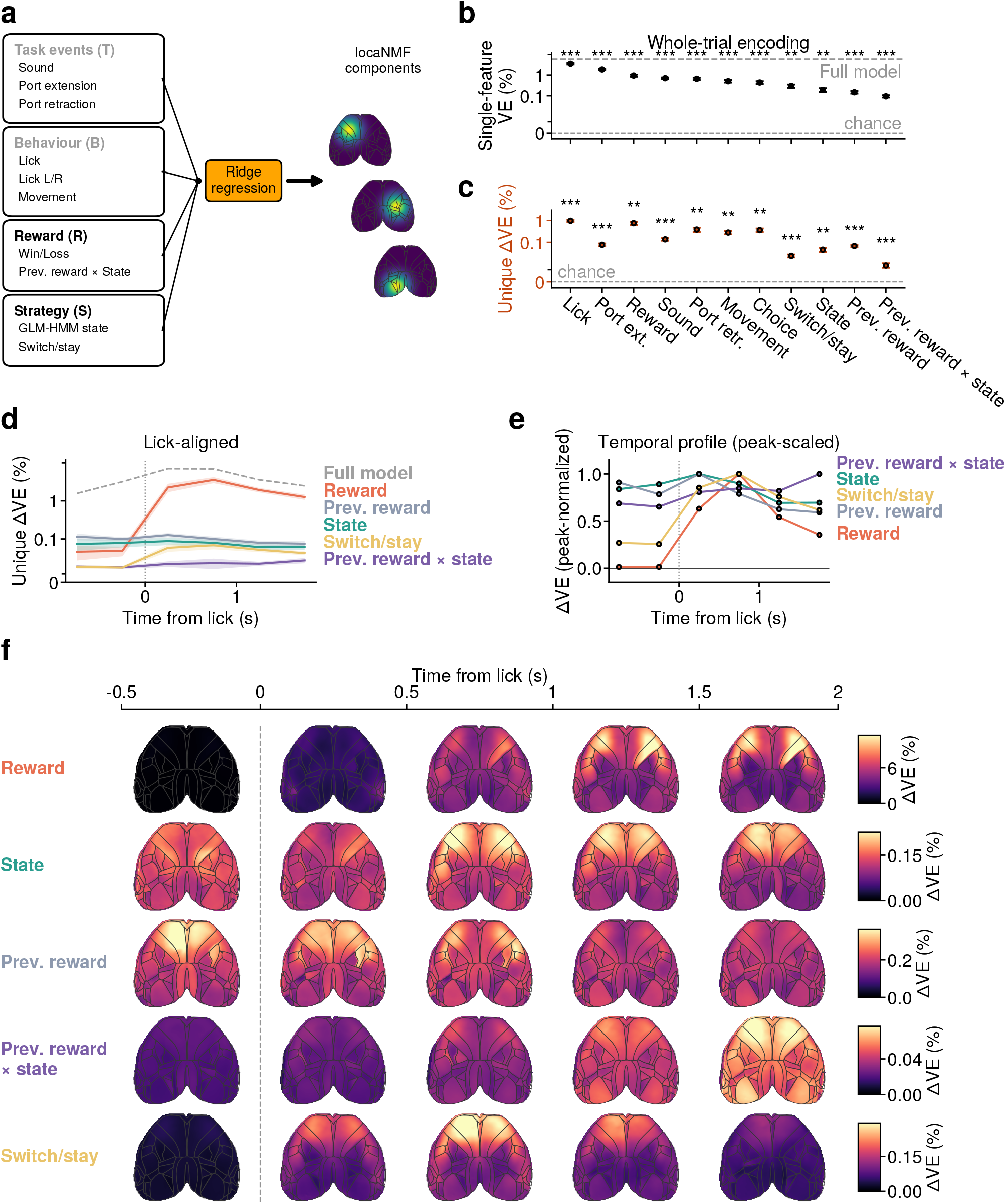
Encoding model of widefield signals during the matching pennies task. **a**, Schematic of the regression model, with the input regressors on the left, and the target output being the locaNMF components. **b**, Variance explained using each of the regressors in a separate encoding model. **c**, Unique variance explained by each regressor in the full encoding model. **d**, Lick-aligned unique variance explained by regressors. **e**, Same as d but scaled to the peak. f, Widefield maps showing the spatial map of unique variance explained by regressors, obtained by scaling the spatial weights of the locaNMF components by their variance explained.

**Extended Figure 9.**
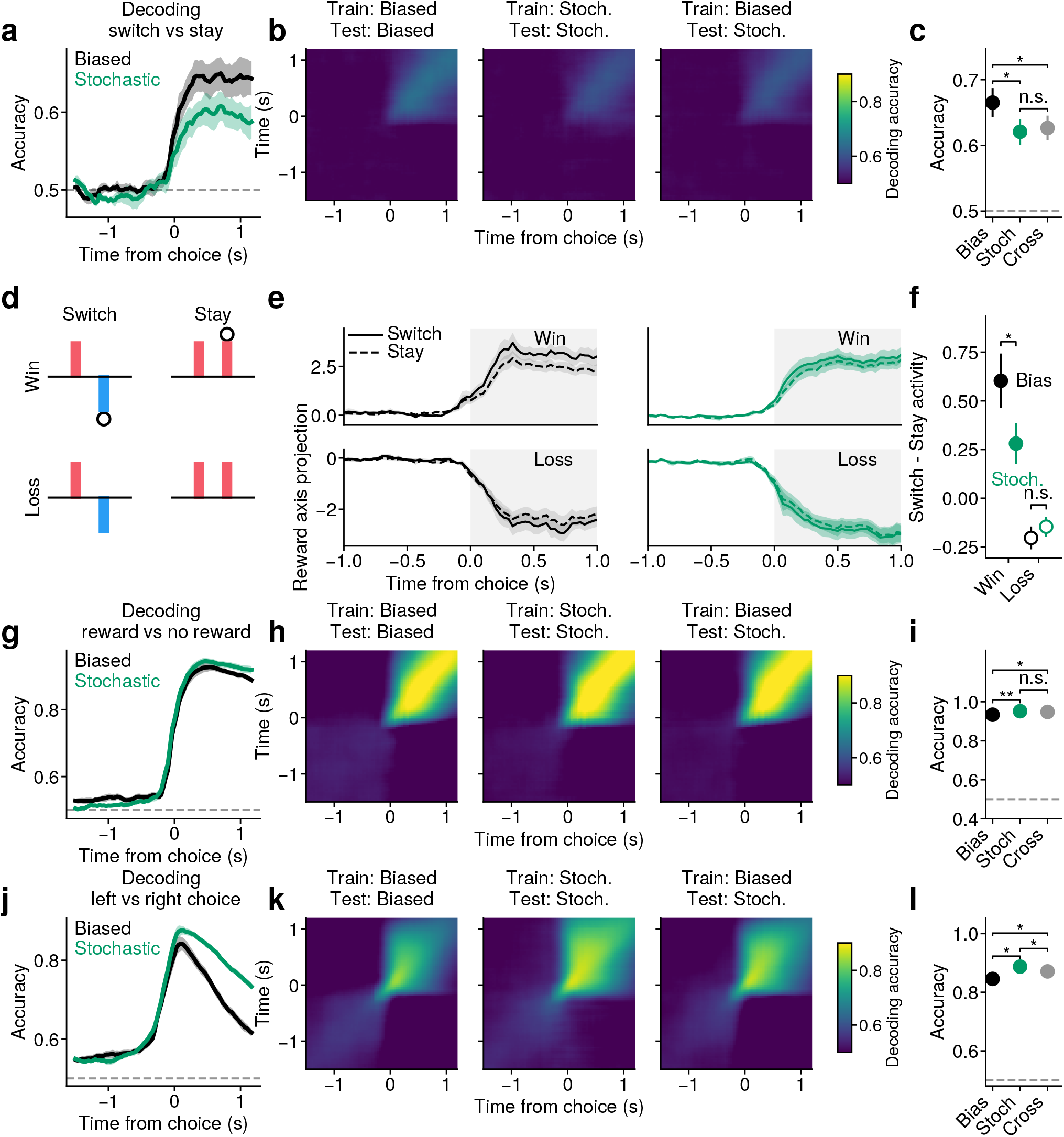
Switch/stay, reward and choice signals in widefield activity. **a**, Decoding accuracy of switch versus stay in biased and stochastic states, a separate decoder is fit to each time window. **b**, Left: Cross-window decoding accuracy within biased states for switch/stay decoding. Middle: Same but for stochastic states. Right: Cross-window decoding accuracy where decoder was trained on biased states and tested on stochastic states. **c**, Statistical comparison of peak switch/stay decoding accuracy within biased, within stochastic and across states (n = 7 mice, paired t-test). **d**, Schematic of the four trial types comparing switch and stay trials conditioned on current reward. **e**, Left: In the biased state, activity projected onto the current reward decoding axis during rewarded (top row) and unrewarded (bottom row) trials for switch (solid line) and stay choices (dashed line). Middle: Same as left but in the stochastic state. Right: Difference in activity between switch versus stay during biased (black) and stochastic (green) state. f, Difference in switch versus stay activity during biased and stochastic state for rewarded and unrewarded trials (n = 7 mice, paired t-test). **g**, Same as a but for reward decoding. **h**, Same as b but for reward decoding. **i**, Same as c but for reward decoding. **j**, Same as a but for choice decoding. **k**, Same as b but for choice decoding. **l**, Same as c but for choice decoding. * p < 0.05, ** p < 0.01, *** p < 0.001, **** p < 0.0001, n.s. not significant.

## Notes

### Competing Interest Statement

The authors have declared no competing interest.

### Summary of Updates

Improved clarity on text and figures.

